# Synthetase and Hydrolase Specificity Collectively Excludes 2’-Deoxyguanosine from Bacterial Alarmone

**DOI:** 10.1101/2024.01.06.574488

**Authors:** Rich W. Zhou, Isis J. Gordon, Yuanyuan Hei, Boyuan Wang

## Abstract

In response to starvation, virtually all bacteria pyrophosphorylate the 3’-hydroxy group of GTP or GDP to produce two messenger nucleotides collectively denoted as (p)ppGpp. Also known as alarmones, (p)ppGpp reprograms bacterial physiology to arrest growth and promote survival. Intriguingly, although cellular concentration of dGTP is two orders of magnitude lower than that of GTP, alarmone synthetases are highly selective against using 2’-deoxyguanosine (2dG) nucleotides as substrates. We thus hypothesize that production of 2dG alarmone, (p)pp(dG)pp, is highly deleterious, which drives a strong negative selection to exclude 2dG nucleotides from alarmone signaling. In this work, we show that the *B. subtilis* SasB synthetase prefers GDP over dGDP with 65,000-fold higher *k_cat_/K_m_*, a specificity stricter than RNA polymerase selecting against 2’-deoxynucleotides. Using comparative chemical proteomics, we found that although most known alarmone-binding proteins in *Escherichia coli* cannot distinguish ppGpp from pp(dG)pp, hydrolysis of pp(dG)pp by the essential hydrolase, SpoT, is 1,000-fold slower. This inability to degrade 2’-deoxy-3’-pyrophosphorylated substrate is a common feature of the alarmone hydrolase family. We further show that SpoT is a binuclear metallopyrophoshohydrolase and that hydrolysis of ppGpp and pp(dG)pp shares the same metal dependence. Our results support a model in which 2’-OH directly coordinates the Mn^2+^ at SpoT active center to stabilize the hydrolysis-productive conformation of ppGpp. Taken together, our study reveals a vital role of 2’-OH in alarmone degradation, provides new insight on the catalytic mechanism of alarmone hydrolases, and leads to the conclusion that 2dG nucleotides must be strictly excluded from alarmone synthesis because bacteria lack the key machinery to down-regulate such products.

## Introduction

Hyperphosphorylated guanosine nucleotides are an important class of bacterial second messenger ^1-3^. Produced by transferring the β,γ-diphosphate of ATP to the 3’-OH of GTP or GDP, these nucleotides are collectively denoted as (p)ppGpp (Figure 1A). First discovered over half a century ago in amino-acid starved *E. coli*, these nucleotides are often referred to as “alarmones” for their functions to alert bacteria to starvation and to reprogram cells to arrest growth and promote survival ^4^. This process, known as the stringent response, is key to the fitness of bacteria under fluctuating conditions and plays a pivotal role in pathogenesis of many infectious species^5^. *E. coli* encodes two alarmone synthetases, RelA and SpoT ^6, 7^(Figure 1A). SpoT also possesses a hydrolase domain that cleaves the 3’-diphosphate from alarmones ^6^. RelA shares the same domain architecture to SpoT, but RelA hydrolase domain is a pseudoenzyme bearing inactivating mutations. Nevertheless, both alarmone synthetases and hydrolases are termed *<u>R</u>*elA-*<u>S</u>*poT *<u>h</u>*omolog (RSH) proteins ^8^. Multiple mechanisms have been reported to trigger alarmone accumulation in response to starvation. For example, RelA associates to translating ribosomes and is activated by uncharged tRNA as a signal of amino acid-starvation ^9^. SpoT senses signals including *apo* acyl carrier protein (fatty-acid starvation), YtfK protein (carbon starvation) and the lack of branched-chain amino acids in favor of alarmone acculumation^10-12^. *E. coli* also encodes GppA, a hydrolase that converts pppGpp to ppGpp, making ppGpp the predominant alarmone species in *E. coli* (Figure 1A, top) ^13^. Once accumulated in bacterial cells, (p)ppGpp bind many protein effectors to modulate their activity, stability, or interactions ^3, 14, 15^. In *E. coli*, for example, ppGpp reprograms transcription by directly binding to RNA polymerase, down-regulating translation by inhibiting multiple ribosome-associated GTPases and modulating nucleotide and amino-acid metabolism by interacting with related metabolic enzymes. Despite efforts to identify (p)ppGpp binding proteins from several bacterial organisms^16-20^, how (p)ppGpp tips the balance between growth and survival is not completely understood.

**Figure 1.**
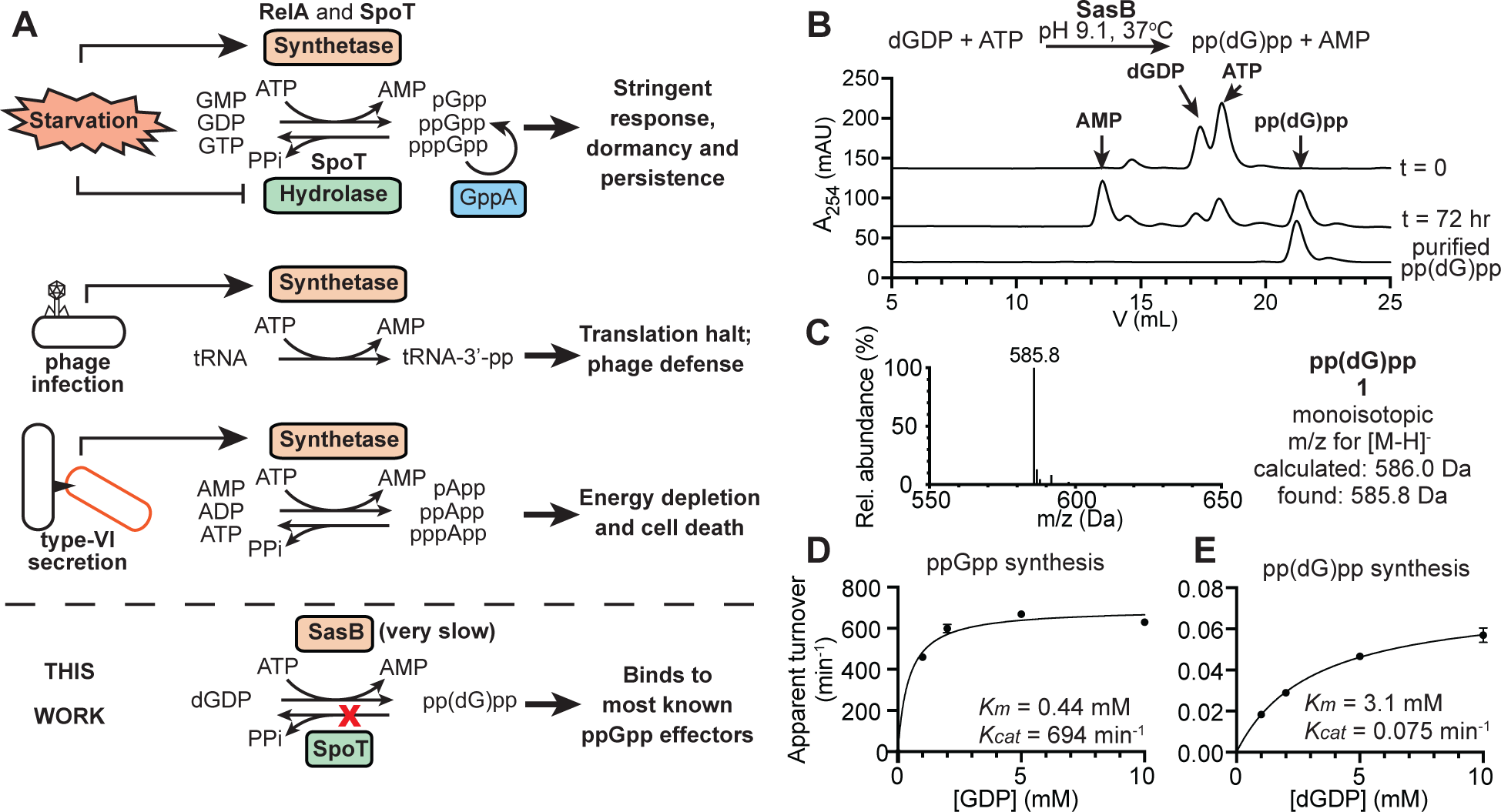
pp(dG)pp synthetase activity of *B. subtilis* SasB. (**A**) Known biochemical activities and functions of RSH synthetases. (**B**) Anion-exchange chromatogram showing the progression of pp(dG)pp synthesis by SasB. Arrows indicate elution volume of standard nucleotides; (**C**) ESI-MS of pp(dG)pp in negative mode. The compound was precipitated from concentrated anion-exchange fractions using LiCl and ethanol and then re-dissolved in water prior to analysis. (**D** and **E**) Kinetics of ppGpp (**D**) and pp(dG)pp (**E**) synthesis as a function of [GDP] or [dGDP]. Fitting curve for a Michaelis-Menten model and parameters are shown. All reactions were carried out in the presence of 10 mM ATP. Apparent turnover is calculated as the initial velocity in μM/min divided by SasB concentration in μM. Error bar represents the range of two reactions set up separately.

Recently, several RSH synthetases were found to pyrophosphorylate the 3’-end of transfer RNAs and 3’-OH of adenosine nucleotides ^21, 22^. Their products are no longer capable of supporting protein translation and energy metabolism, respectively (Figure 1A). Activities of these homologs are so strong that they cause cytotoxicity. Genes of these RSH-synthetase toxins are found co-operonic with their antitoxin genes encoding an antidote to ensure survival of the host ^23^. Activation of these toxins occurs through either recognizing phage proteins in the host or being secreted into another bacterial cell ^21, 24^. In the first scenario, the toxicity sacrifices the infected host cell to abort phage replication and protect the bacterial population while in the second, the toxin antagonizes the growth of the recipient cell, potentially from a competing species, to establish the host’s advantage in the niche ^21, 24^ (Figure 1A). These new discoveries show that RSH synthetases can evolve to adapt pyrophosphate-accepting substrates with different nucleobase and/or 5’-appendages. Nonetheless, no RSH synthetase has been observed to tolerate 2’-deoxy modification of the substrate. Recently, Patil et al. showed that dGTP binds alarmone synthetases with similar affinity to GTP but is not a substrate ^25^, suggesting a negative-selection pressure that excludes 2dG nucleotides from alarmone signaling. Furthermore, because cellular concentrations of dGTP (ca. 50 μM) are 2 orders of magnitude lower than GTP (ca. 5 mM) ^26, 27^, an alarmone synthetase lacking specificity against 2dG substrates would only produce a small quantity of (p)pp(dG)pp. We therefore hypothesize that production of (p)pp(dG)pp is highly deleterious, which drives the strong ribonucleotide specificity of RSH synthetases.

To test our hypothesis, we first quantified the GDP versus dGDP specificity of *B. subtilis* SasB (also known as YjbM, RelQ or SAS1), a single-domain RSH synthetase. SasB utililzes dGDP with 65,000-fold lower *k_cat_*/*K_m_*, a specificity to exclude 2’-deoxynucleotides stricter than RNA polymerases. To explore the potential fitness consequence of (p)pp(dG)pp production, we used chemical proteomics to identify *E. coli* proteins binding to pp(dG)pp or ppGpp, which led to the surprising discovery that pp(dG)pp is a poor substrate of ppGpp hydrolase, SpoT (Figure 1A, bottom). This inability to hydrolyze 2’-deoxy hyperphosphorylated substrate is conserved across several distantly related RSH hydrolyases. Our biochemical data show that 2’-OH plays a key role in stabilizing the catalytically productive conformation of SpoT-ppGpp complex, likely by direct coordination to the Mn^2+^ at the SpoT catalytic center. Our work revealed a previously overlooked, vital role of 2’-OH in alarmone degradation, provided new insights to the catalytic mechanisms of RSH hydrolases and opens the possibility that the inability to hydrolyze pp(dG)pp constrains the nucleotide specificity of alarmone signaling.

## Results

### *B. subtilis* SasB has a slow pp(dG)pp-synthetase activity

*B. subtilis* SasB is a single-domain RSH synthetase allosterically activated by pppGpp^28^. Given its stability and high activity in the activated state, SasB is widely used for (p)ppGpp synthesis *in vitro*. Importantly, Patil et al. reported that SasB shows stronger activity at pH 8.5 than at 6.5 when pyrophosphorylating GTP analogs in which 2’-OH is replaced with a fluorine or an amino group^25^. We thus used high pH (9.1) and a high concentration of SasB (50 μM) to drive the difficult synthesis of pp(dG)pp. In 72 hours, a majority of dGDP was hyperphosphorylated (Figure 1B), which was confirmed to be pp(dG)pp (**1**) using MS (Figure 1C) and NMR (SI-NMR document). Surprised by this new activity, we characterized its Michaelis-Menten kinetics at pH 7.8 and 30°C. Synthesis of pp(dG)pp by SasB has a *k_cat_* of 0.075/min and *K_m_* of 3.1 mM for dGDP, compared to 694/min and 0.44 mM for GDP in ppGpp synthesis (Figure 1D and E). These constitute a 65,000-fold difference in *k_cat_/K_m_* in favor of GDP as the diphosphate acceptor. The specificity is much stricter than the selection of rNTPs for transcription elongation by bacterial, eukaryotic and viral RNA polymerases, which typically come with a *k_cat_/K_m_* difference between 100-1,000 folds ^29-31^, indicating a strong selection pressure against the synthesis of pp(dG)pp.

### pp(dG)pp and ppGpp have near-identical sets of binding proteins

Because of the extremely weak pp(dG)pp synthetase activity and the direct competition between GDP and dGDP for the active center^25^, expressing SasB or other ppGpp synthetases in bacterial cells is unlikely to produce a substantial amount of pp(dG)pp. To understand the potential consequence of pp(dG)pp production were it ever to get made at a significant level, we instead use capture compound mass spectrometry (CCMS) to compare the binding of ppGpp and pp(dG)pp to proteins in *E. coli* lysate (Figure 2A). A similar approach has been used by us and others to identify dozens of ppGpp-binding proteins in *E. coli* ^19, 20^. Given the structural similarity between pp(dG)pp and ppGpp, we expect that they should share most binding proteins. Nevertheless, if pp(dG)pp is indeed toxic as we hypothesized, it likely acts through a protein effector not shared with ppGpp because ppGpp induction alone does not compromise viability.

**Figure 2.**
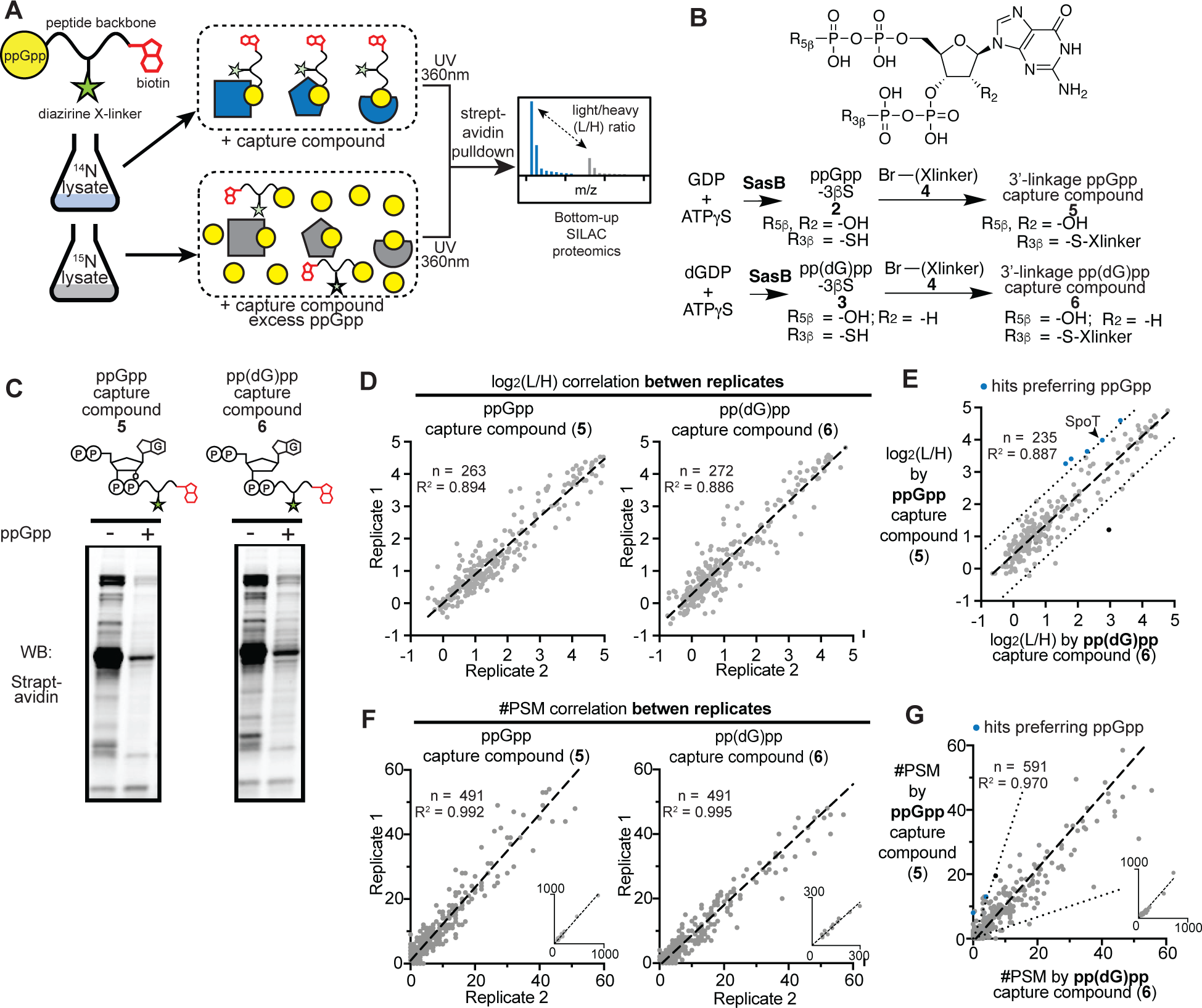
Comparative CCMS between ppGpp and pp(dG)pp capture compounds. (**A**) CCMS approach using stable isotope-labeled bacterial lysates. (**B**) Scheme showing the synthesis of ppGpp and pp(dG)pp capture compounds. See **SI Figure 1** for structure and LC-MS characterization of the bromoacetylated crosslinker peptide (**4**). (**C**) Tagged *E. coli* proteome by indicated capture compounds visualized using streptavidin western blotting. *E. coli* lysate containing 200 μM capture compound (-ppGpp) or 200μM capture compound and 5 mM ppGpp (+ ppGpp) was illuminated with 360-nm UV lamp. Aliquots containing 15 μg protein was resolved on 12% SDS-PAGE for immunoblotting. (**C**) Statistics of peptides and proteins detected in six CCMS datasets. (**D** and **E**) Log_2_(L/H) correlation analysis between replicates (panel **D**) and between 3’-linkage ppGpp versus pp(dG)pp capture compounds (panel **E**). Each dot represents the Log_2_(L/H) value of a protein reliably detected (see **Table 1** for criteria) in both replicates. Regression line is shown in dash. In (**E**), the mean Log_2_(L/H) value from two replicates was used for plotting. Dotted lines are one Log_2_ unit above or below the regression line and set the threshold for differentially captured hits favoring 3’-linkage ppGpp (blue dots) (see **Table 1**). The black data point represents Gsk, a hit showing opposite preference in #PSM analysis. (**F** and **G**) Same to (**D** and **E**), except #PSM of all proteins detected by as few as one peptide was plotted for analysis. Each panel is a zoom-in for #PSM ≤ 60 with an inset showing all data points outside.

**Table 1.** Hit proteins preferentially crosslinked to ppGpp capture compound.

| Uniprot Accession | Gene | Protein description | Hit by capture compound | $\log_2(H/L)$ difference* | # PSMs ratio* |
| --- | --- | --- | --- | --- | --- |
| P0AG24 | *spoT* | Bifunctional (p)ppGpp synthase/hydrolase SpoT | both | 1.22 | 3.0 |
| P00962 | *glnS* | Glutamine--tRNA ligase | both | 1.35 | 1.4 |
| P46853 | *yhhX* | Uncharacterized oxidoreductase YhhX | both | 1.28 | 1.6 |
| P0A6T5 | *folE* | GTP cyclohydrolase 1 | ppGpp only | 1.61 | 2.3 |
| P0A6P1 | *tsf* | Elongation factor Ts | ppGpp only | 1.64 | 1.8 |
| P0A6A3 | *ackA* | Acetate kinase | ppGpp only | NA | 3.3 |
| P0A6A4 | *pssA* | CDP-diacylglycerol--serine O-phosphatidyltransferase | ppGpp only | NA | NA |

\* NA occurs when no $\log_2(L/H)$ or PSM values were found using pp(dG)pp probe.

To identify effectors proteins that can distinguish between ppGpp and pp(dG)pp, we generated photo-crosslinking probes for both nucleotides, hereafter referred to as “capture compounds”, to carry out a comparative CCMS study (Figure 2B). We first used SasB to synthesize analogs of ppGpp and pp(dG)pp in which the 3’-β-phosphate is substituted by a thiophosphate (Figure 2B). These nucleotides, G4P3βS (**2**) and dG4P3βS (**3**), were then conjugated to a bromoacetylated peptide bearing a diazirine moiety and a biotin handle (**4,** see also SI Figure 1) to give rise capture compounds **5** and **6**. We then used each capture compound to covalently tag its binding proteins in a *E. coli* lysate for enrichment and identification using MS. To enable quantitative MS readout, we grew *E. coli* in M9-glucose (M9G) minimal medium using light (^14^N) or heavy (^15^N) ammonium chloride as the sole nitrogen source (Figure 2A). Cells were harvested in mid-exponential phase and lysed. We performed a pair of crosslinking reactions for each capture compound: we added 100 μM each capture compound to both lysates, and excess, 5.0 mM native ppGpp to only the heavy lysate to compete with the capture compound for binding. After UV exposure, covalent crosslinking effectively biotinylates proteins bound to capture compounds. Notably, both ppGpp and pp(dG)pp capture compounds produced similar pattern of biotinylated proteins on a streptavidin immunoblot (Figure 2C, compare left lanes), in agreement to our expectation of their overlapping interactomes. Presence of native ppGpp strongly diminished protein tagging by all capture compounds (Figure 2C, compare two lanes within each blot), suggesting that most tagged proteins bind ppGpp.

We next combined crosslinking reactions with and without excess ppGpp for each capture compound and used immobilized streptavidin to enrich biotinylated proteins, which were then digested for LC-MS analysis (Figure 2A). Each capture compound was tested in two biological replicates using *E. coli* lysates prepared from separate cultures. Tryptic fragments from ppGpp or pp(dG)pp-binding proteins are expected to be enriched in ^14^N isotope. Thus, we use the average abundance ratio of ^14^N over ^15^N tryptic fragments of each protein, denoted as ‘L/H’ hereafter, as a metric of the protein’s interaction specificity to the nucleotide-of-interest. For proteins detected by at least two unique tryptic fragments in both replicates, their log_2_(L/H) correlates well between replicates, suggesting excellent reproducibility (Figure 2D). Remarkably, for proteins identified by both ppGpp and pp(dG)pp capture compounds, their log_2_(L/H) correlates equally well across capture compounds with a squared Pearson correlation coefficient, R^2^ of 0.887 compared to 0.894 and 0.886 between replicates (Figure 2E). Since proteins identified only in one dataset would be excluded from Log_2_(L/H) correlation analysis, we also used the total number of *<u>p</u>*eptide-*<u>s</u>*pectrum *<u>m</u>*atches (#PSM) as a metric of the crosslinking yield for each protein. Strong correlation of #PSM was also observed between capture compounds and the R^2^ of 0.970 was only slightly lower than 0.992 and 0.995 found between replicates (Figure 2F and G). For each capture compound, we selected top 20% protein of highest L/H as hits for ppGpp- or pp(dG)pp binding, leading to 55 and 56 hits for ppGpp and pp(dG)pp capture compound, respectively, with an intersection of 45 (SI Spreadsheet 1).

To provide a quantitative view on the sensitivity of ppGpp-binding protein to the missing 2’-OH in pp(dG)pp, we selected four *E. coli* ppGpp effectors, namely constitutive ornithine decarboxylase (SpeC), inducible lysine decarboxylase (LdcI), inosine-guanosine kinase (Gsk) and amidophosphoribosyltransferase (PurF) and compared their binding enthalpy and affinity with ppGpp and pp(dG)pp using isothermal titration calorimetry (ITC). In published ppGpp-bound crystal structures, 2’-OH makes no contact with LdcI and accepts long (> 3.5 Å) H-bonds from Gsk and PurF ^19, 32, 33^. Nevertheless, affinity of pp(dG)pp to all for proteins are on the same order to that of ppGpp (SI Figure 2). Taken together, interactomes of ppGpp and pp(dG)pp largely overlap, suggesting that 2’-OH is not a requirement for ppGpp binding to most protein effectors.

### 2’-OH is required for the activity of RSH hydrolases

Since the hypothesized detrimental effect of pp(dG)pp requires one or more effector proteins that responds differently to ppGpp and pp(dG)pp, we examined hits identified with most different Log_2_(L/H) or #PSMs by two capture compounds. Table 1 summarized 7 differentially captured hits (Figure 2E and F, blue symbols) selected using following thresholds (Figure 2E and F, dotted lines): either their Log_2_(L/H) found by the pp(dG)pp capture compound deviates from the regression line by more than one Log_2_ unit or their #PSM differs by greater than 3 folds between capture compounds. All 7 hits were identified with stronger crosslinking to ppGpp capture compounds, among which SpoT is the only known ppGpp-binding protein (Figure 2E, arrowhead). Since there is no obvious structural or functional commonplace shared by these hits, we next focused on validating whether SpoT can discern between ppGpp and pp(dG)pp. *E. coli* SpoT is an essential protein because it possesses the only hydrolase activity that effectively downregulates alarmones. To test pp(dG)pp hydrolysis by SpoT, we purified full-length recombinant SpoT from *Acinetobacter baumannii*, SpoT*^Ab^*, a homolog that displays much better solution behavior^34^ (See Figure 3A for sequence alignment). Harnessing the activity of nucleotide disphosphate kinase (Ndk), pyruvate kinase (PK) and lactose dehydrogenase (LDH), we coupled the production of GDP or dGDP by SpoT activity to the reduction of NADH, which was detected as a decrease of absorbance at 340 nm (Figure 3B). We compared hydrolysis kinetics for ppGpp and pp(dG)pp by SpoT*^Ab^* at pH 7.5 in the presence of 5 mM MgCl_2_. Strikingly, the missing 2’-OH decreases the *k_cat_* by 100 folds and increases *K_m_* by 9.6 folds (Figure 3C and D), resulting in a 1,000-fold loss in *k*_cat_/*K*_m_. Given the 55% sequence identity between hydrolase domains of SpoT*^Ab^* and *E. coli* SpoT*^Ec^* (Figure 3A), the latter likely shares similar deficiency in pp(dG)pp hydrolysis.

**Figure 3.**
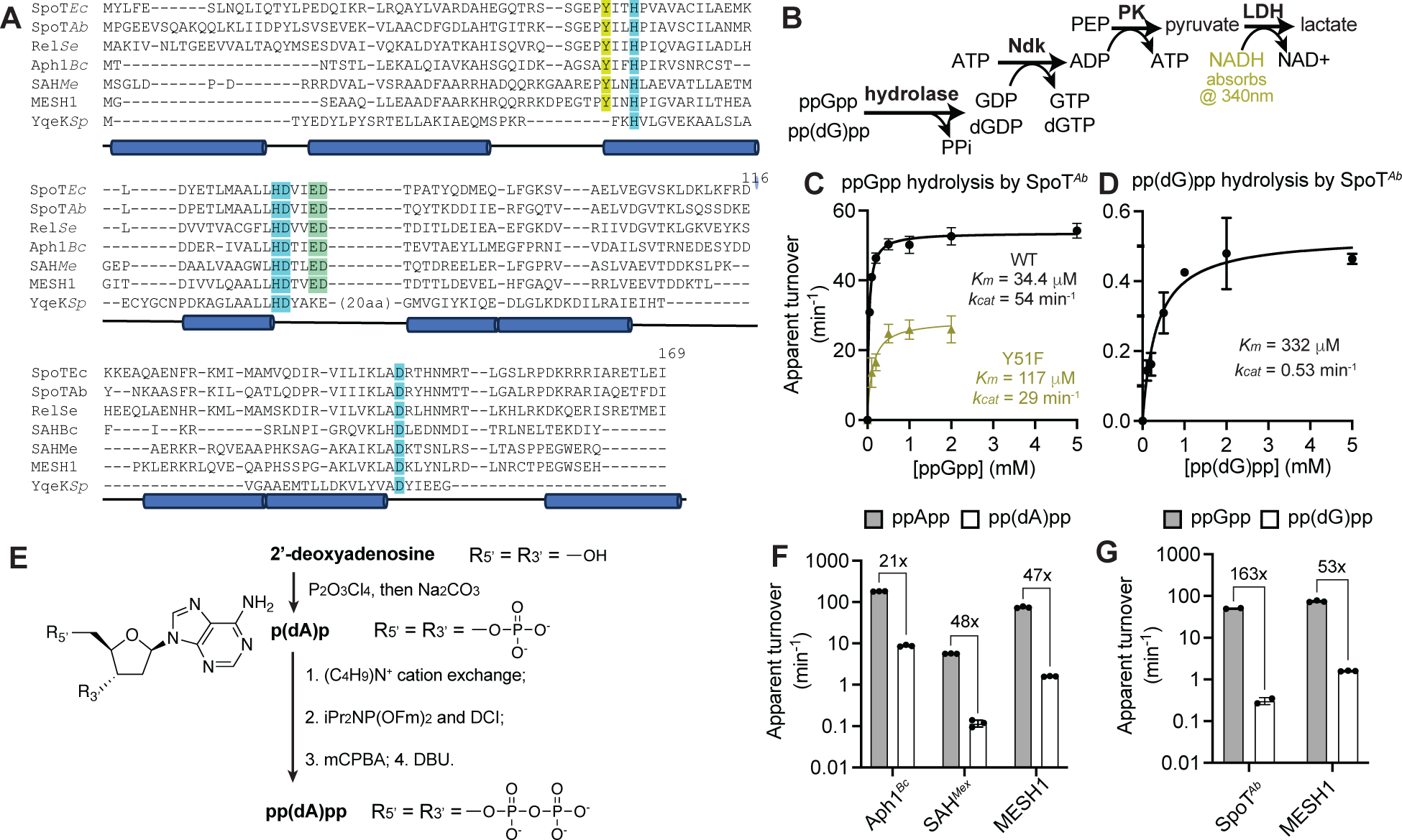
RSH hydrolases have poor activity on 2’-deoxy substrates. (**A**) Sequence alignment of hydrolase domains in long RSH proteins (*E. coli* and *A. baumanii* SpoT and *S. equorum* Rel), SAHs from *B. cereus*, *M. extorquens* and *H. sapiens* and a non-RSH family HD domain hydrolase, *S. pyogenes* YqeK. Numbering of amino-acid positions of SpoT*^Ab^* are used. (**B**) Coupling of RSH hydrolase activity to NADH consumption. (**C** and **D**) Hydrolysis kinetics of ppGpp (**C**) and pp(dG)pp (**D**) by SpoT^Ab^ plotted as functions of substrate concentration and fitted for a Michaelis-Menten model. Error bars indicate range of two reactions assembled separately. (**E**) Scheme showing the synthesis of pp(dA)pp. (**F** and **G**) Activity comparison of indicated hydrolase on 500 μM canonical and 2’-deoxy substrates. To provide appropriate level of absorbance signal, all 2’-deoxy substrate hydrolysis was carried out at 10x enzyme concentration relative to hydrolysis of the canonical substrate. In Panels **C**, **D**, **F** and **G**, apparent turnover is calculated as the initial velocity in μM/min divided by enzyme concentration in μM.

Because 2’-deoxy alarmones have never been detected in any bacterial organism, there is no obvious selection pressure that drives RSH hydrolases to avoid pp(dG)pp. We therefore speculate that the slow hydrolysis of pp(dG)pp may reflect a critical role of 2’-OH in the enzymology shared by all RSH hydrolases. In addition to long RSH proteins possessing both synthetase and hydrolase domains, RSH hydrolases also exist as small, single-domain enzymes known as *<u>s</u>*mall *<u>a</u>*larmone *<u>h</u>*ydrolases (SAHs) (See Figure 3A). To date, most SAHs characterized *in vitro* hydrolyze (pp)pApp, although some bifunctional SAHs also accept (pp)pGpp as substrates ^35-38^. To test SAHs’ activity on 2’-deoxy substrates, we first chemically synthesized pp(dA)pp from 2’-deoxyadenosine by two rounds of 3’,5’-bisphosphorylation using first diphosphoryl-chloride and then phosphoramidate chemistry (Figure 3E) ^39^. In agreement with our speculation, two (pp)pApp-specific SAHs from *Bacteroides caccae* (Aph1*^Bc^*) and *Methylobacterium extorquens* (SAH*^Me^*) and the bifunctional SAHs from *Homo Sapiens* (MESH1) all hydrolyze pp(dA)pp albeit by 21-48 folds slower compared to ppApp (Figure 3F). Similarly, Mesh1 is 53-fold slower in hydrolyzing pp(dG)pp than ppGpp (Figure 3G). Taken together, inefficiency of hydrolyzing 2’-deoxy-3’-pyrophosphorylated nucleotides is probably a common feature of all RSH hydrolases.

### Activity of SpoT*^Ab^* requires two divalent metal cations

To further understand the role of 2’-OH in alarmone hydrolysis, we inspected the ppGpp-bound crystal structures of SpoT*^Ab^* (PDBID: 7QPR and Figure 4A) ^32^. The first and foremost contact involving 2’-OH is a 3.0-Å H-bond to Tyr51, a residue invariant across the RSH hydrolase family (Figure 5A, Tyr51 in yellow sticks and H-bond in black dashes)^8^. Surprisingly, in contrast to the profound effect of 2’-deoxy modification, disrupting this H-bond by the Y51F mutation only increased the *K_m_* by 2.4 fold and decreased the *k_cat_* by 47% for ppGpp hydrolysis (Figure 3C, yellow data points). This result suggests that the 2’-OH has another more significant role in enabling catalysis by SpoT. All RSH hydrolases device four invariant residues, two histidines and two aspartates, hereafter referred to as the 2H2D motif, to accommodate a Mn^2+^ cation^8^. In the SpoT^Ab^-ppGpp complex, this Mn^2+^ also receives coordination from the 3’-α-phosphate (Figures 3A and 4A, 2H2D motif highlighted in light blue and coordination bonds in red dashes). Intriguingly, in all four SpoT^Ab^ subunits in the asymmetric unit, the distance between the 2’-oxygen and this Mn^2+^ is 3.5-4.0 Å (Figure 4A, right panel, green dash). Similar positioning between ppGpp and the Mn^2+^ center is also observed in the crystal structure of another long-RSH protein, Rel from *Thermus thermophilus* (PDBID: 6S2T) ^40^. Notably, both crystal structures were obtained using the wild-type hydrolase and thus the ppGpp observed must deviate from the optimal conformation for hydrolysis (otherwise the nucleotide would have been hydrolyzed). We therefore speculate that the 2’-OH may be in closer and stronger contact with the Mn^2+^ center during catalysis.

**Figure 4.**
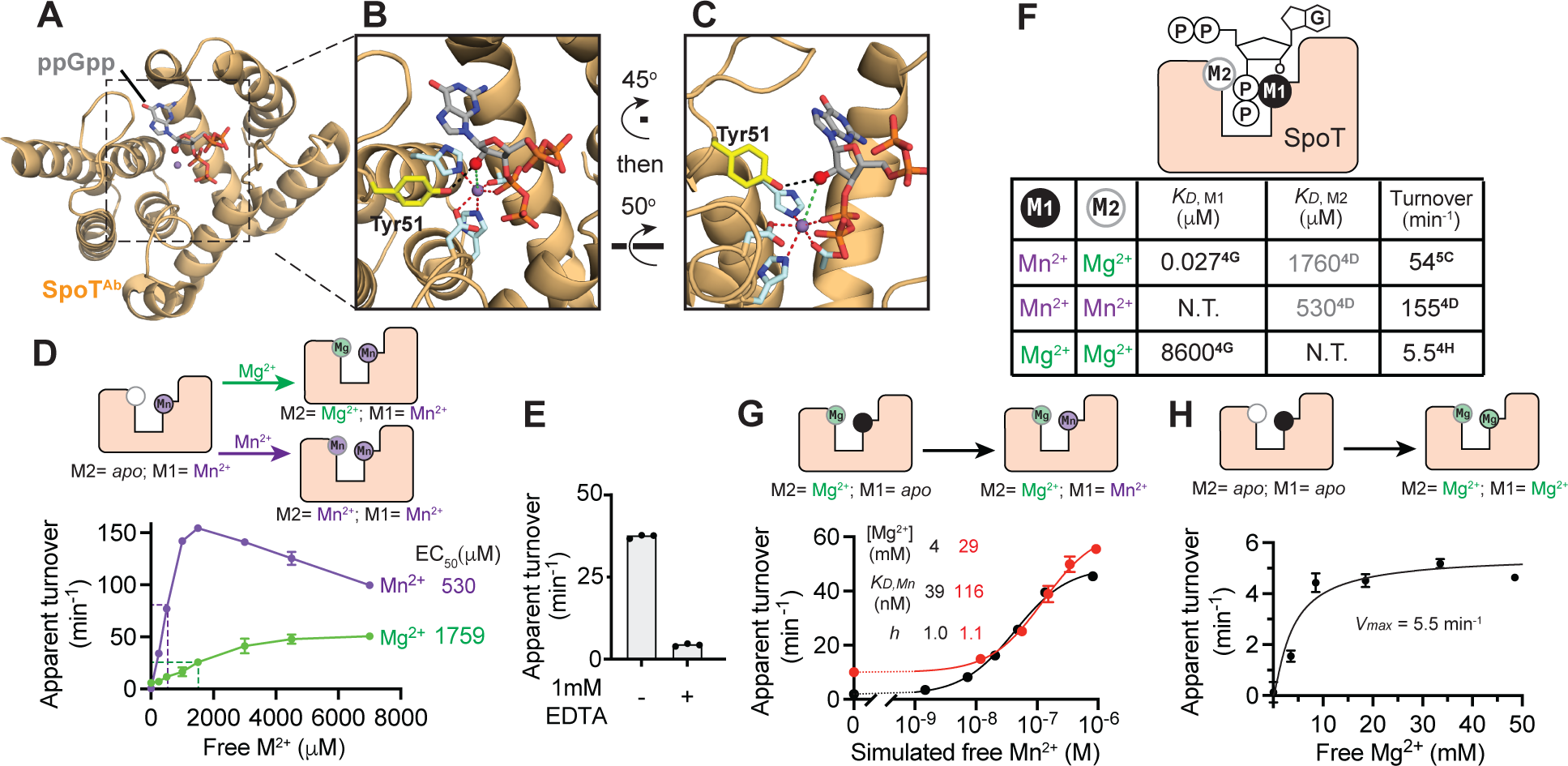
SpoT*^Ab^* is a binuclear metallopyrophosphohydroase. (**A-C**) Overview (**A**) and active-center zoom-in (**B** and **C**) of the published crystal structure of SpoT*^Ab^*-ppGpp complex. ppGpp and side chains of conserved residues related to this study are in stick model. 2’-oxygen is a red sphere and Mn^2+^ is a purple sphere. (**D** and **E**) ppGpp hydrolysis activity of SpoT*^Ab^* as a function of free Mg^2+^ or Mn^2+^ concentration (**D**) and in the presence of 10 mM MgCl_2_ with or without 1 mM EDTA (**E**). Error bars in (**D**) indicate S.D. of three reactions assembled separately. (**F**) Estimated affinity of metal binding and resulted specific activity of SpoT^Ab^ in three different metal-binding states. Superscript in the table indicate the Figure panel from which the parameter is derived. *K_D_* values estimated from EC_50_ are in gray. (**G** and **H**) ppGpp hydrolysis activity of SpoT*^Ab^* as a function as a function of free Mn^2+^ concentration in the presence 10 (black) or 30 mM (red) MgCl_2_ and 1 mM EDTA (**G**) and as a function of free Mg^2+^ concentration in the presence of 1 mM EDTA (**H**). Error bars indicate range of two reactions assembled separately. Data were fitted to Hill equation in (**G**) and a model in which activity requires two Mg^2+^ binding independently at respective *K_D_* of 1.76 and 8.6 mM in (**H**). In panels **D**, **E**, **G** and **H**, apparent turnover is calculated as the initial velocity in μM/min divided by enzyme concentration in μM.

**Figure 5.**
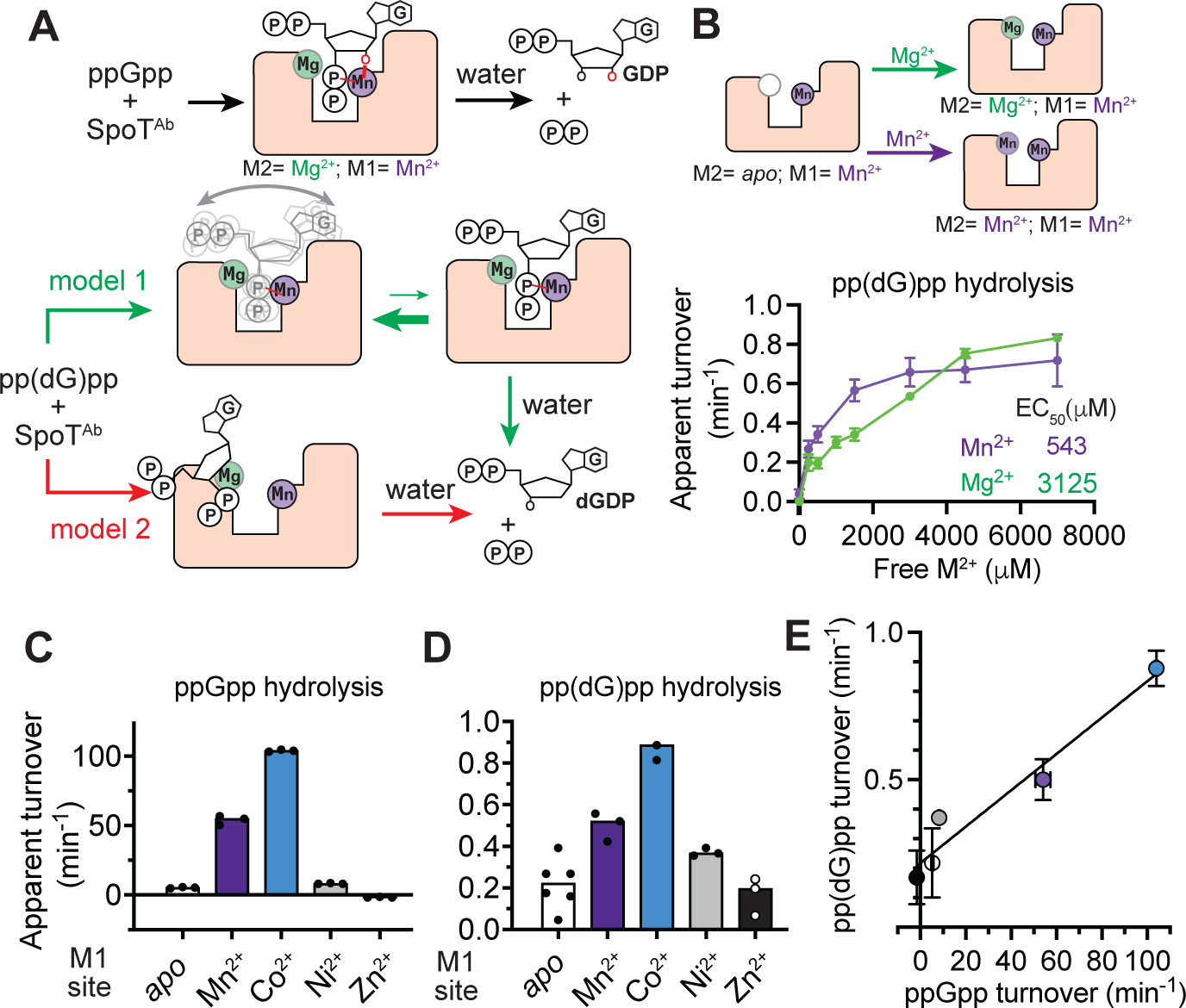
ppGpp and pp(dG)pp hydrolysis share similar metal dependence for both M1 and M2 sites. (**A**) Two models accounting for the slow hydrolysis of pp(dG)pp. pp(dG)pp either has a much lower chance to adopt to the same conformation required for ppGpp hydrolysis (green arrows) or goes through a different transition state (red arrow). (**B**) ppGpp hydrolysis activity of SpoT*^Ab^* as a function of free Mg^2+^ or Mn^2+^ concentration. Error bars indicate S.D. among three reactions assembled separately. (**C** and **D**) Hydrolysis kinetics of ppGpp (**C**) and (**D**) in the presence of 10mM MgCl_2_ and 100 μM indicated metal that saturates the M1 site. 1 mM EDTA was included in “*apo*” reactions. (**E**) Correlation plot of pp(dG)pp and ppGpp hydrolysis activities in panels **C** and **D** showing the linear relationship when varying the metal cation bound to the M1 site. In Panels **C**-**E**, apparent turnover is calculated as the initial velocity in μM/min divided by enzyme concentration in μM.

Before interrogating the role of this putative 2’-OH-metal interaction, we first clarified the requirements of divalent metal cations for ppGpp hydrolysis by SpoT*^Ab^*. Earlier works have revealed controversial metal-dependence for different RSH hydrolases. For example, a SAH from *Corynebacterium glutamicum*, RelH*^Cg^*, exhibits maximal activity at 1 mM Mn^2+^ or 7.5 mM Mg^2+^, suggesting a low-mM affinity binding site for these cations in the RelH*^Cg^*-ppGpp catalytic complex^36^. We observed similar activation isotherms for SpoT*^Ab^* by Mg^2+^ and Mn^2+^, with half-maximal activity reached at EC_50_ = 1.76 mM free Mg^2+^ or EC_50_ = 530 μM free Mn^2+^ (Figure 4D). It is noteworthy that Mg^2+^ and Mn^2+^ bind ppGpp at 1:1 stoichiometry with 125 μM and 5.5 μM *K_D_*, respectively (SI Figure3 A and B) and their chelation by ppGpp was taken into consideration when we calculated free cation concentrations for Figure 5B. Also similar to RelH*^Cg^* was the decline of hydrolase activity at high Mn^2+^ concentrations by a yet unknown mechanism (Figure 4D, purple trace). In contrast to this low-affinity response to metal cations, SpoT^Ab^ was reported to require exclusively Mn^2+^ at much stronger affinity ^34^. Intriguingly, we also readily reproduced key results in this literature: SpoT*^Ab^* purified using Mn^2+^-free buffer was active in the presence of 10 mM MgCl_2_, while the activity decreased by 90% upon addition of 1mM EDTA (Figure 4E). Importantly, EDTA has 5 orders-of-magnitude stronger affinity to Mn^2+^ than to Mg^2+^ and is hence able to compete Mn^2+^ off the enzyme despite the presence of excess free Mg^2+^. These data conclusively show that (1) SpoT^Ab^ can co-purify with Mn^2+^ and remains active in a Mn^2+^-free buffer, (2) the SpoT^Ab^-Mn^2+^ complex has fast dissociation kinetics to enable inhibition by EDTA; and, therefore, (3) the SpoT^Ab^-Mn^2+^ complex has nanomolar affinity so it can stay thermodynamically stable at a working concentration of 100 nM in our assays. To reconcile this high-affinity Mn^2+^ requirement with the low-affinity dose-response for Mn^2+^ or Mg^2+^ in Figure 4D, we propose that SpoT^Ab^-ppGpp catalytic complex contains two binding sites for divalent metal cations: a high-affinity, M1 site for the Mn^2+^ observed in crystal structure (Figure 4F, black circle) and a previously unreported, low-affinity M2 site for Mg^2+^ or Mn^2+^ (Figure 4F, gray circle). Binding of appropriate cation at both sites are necessary for efficient catalysis. In this model, SpoT^Ab^ activation isotherm in Figure 4D reflects saturation binding of the M2 site provided that M1 site is occupied by Mn^2+^. Mn^2+^ at M2 site provides 2-fold stronger hydrolase activity than Mg^2+^. Assuming that EC_50_ values are a good estimation of *K_D_*, the physiologically relevant ligand of M2 site should be Mg^2+^ given the estimated free-metal concentration on the order of 1 mM for Mg^2+^ (EC_50_ = 1.76 mM) and 1 μM for Mn^2+^ (EC_50_ = 530 μM) ^41^.

To estimate the affinity of Mn^2+^ binding at M1 site, we titrated Mn^2+^ up to 0.9 mM to SpoT^Ab^ hydrolysis reactions in the presence of 5 mM Mg^2+^ and 1 mM EDTA. Under these conditions, dissociation equilibrium of the Mn-EDTA complex controls the free Mn^2+^ at low concentrations resembling conjugate acid-base pair controlling the pH, allowing us to vary free Mn^2+^ concentration from high-picomolar to high-nanomolar ranges (See **Methods** Section for free-Mn^2+^ calculation for each titration point). Fitting the activation isotherm of SpoT*^Ab^* to Hill’s equation provides an apparent *K_D_* = 39 nM and a Hill coefficient = 1.0, in agreement to the well-defined, single Mn^2+^ binding site present in the crystal structure. According to the apparent *K_D_* value, at 100 nM working concentration, SpoT^Ab^-Mn^2+^ complex would dissociate by 45%, consistent with the robust hydrolase activity of our SpoT^Ab^ protein in Mn^2+^-free, EDTA-free buffer. Intriguingly, increasing Mg^2+^ concentration to 30 mM increased the apparent *K_D_* for Mn^2+^ to 116 nM (Figure 4G). We think this increase is due to direct competition between Mg^2+^ and Mn^2+^ for the M1 site, and thus calculated the *K_D_* = 8.6 mM for Mg^2+^ and 29 nM for Mn^2+^, respectively (See **Methods** section for *K_D_* calculation). To test the efficacy of Mg^2+^ binding at M1 site for hydrolase activity, we titrated ppGpp hydrolysis reactions containing 1 mM EDTA with Mg^2+^ and, gratifyingly, restored low level of hydrolase activity with high Mg^2+^ (Figure 4H). Using *K_D_*_, M2_ = 1.76 mM (EC_50_ of Mg^2+^ for M2 site, Figure 4D), and *K_D_*_, M1_ =8.6 mM (calculated *K_D_* of Mg^2+^ for M1), we fitted our data closely to a model in which Mg^2+^ binding at both sites is required for activity. Turnover number with Mg^2+^ binding at both sites is 5.5 min_-1_, about 10-fold lower compared to when M1 is occupied by Mn^2+^. Taken together, our data unambiguously demonstrate that SpoT is a binuclear metallopyrophosphohydrolase.

### pp(dG)pp and ppGpp hydrolysis shares near-identical metal requirement

We next set out to investigate how the 2’-deoxy modification leads to the poor *k_cat_* of pp(dG)pp hydrolysis. Assuming that 2’-OH plays a critical role in docking ppGpp to the M1-site Mn^2+^ as suggested by the SpoT^Ab^-ppGpp crystal structure (Figure 5A, black arrows), the residual activity of pp(dG)pp hydrolysis by SpoT^Ab^ may come from two mechanisms. In the first mechanism, pp(dG)pp is hydrolyzed *via* a transition state analogous to that of ppGpp hydrolysis, while the slow kinetics is caused by the difficulty of pp(dG)pp sampling the productive conformation without the putative 2’-OH-M1 interaction (Figure 5A, green arrows). The second possible mechanism relies on a slow, M1-independent hydrolase activity that is manifested only by pp(dG)pp (Figure 6A, red arrows). Importantly, the first model predicts that hydrolysis rate of ppGpp and pp(dG)pp to respond similarly to the variation of divalent metals binding at both sites in the transition state. In light of this idea, we first varied occupancy of M2 site by carrying out pp(dG)pp hydrolysis at a series of Mg^2+^ or Mn^2+^ concentrations. As shown in Figure 5B, SpoT^Ab^ retains its low-affinity requirement for M2-site cation, with a moderate increase of EC_50_ by 77% for Mg^2+^ and 2% for Mn^2+^ compared to ppGpp hydrolysis. Notably, while Mg^2+^ binds pp(dG)pp and ppGpp indistinguishably in the absence of protein (SI Figure 3C), Mn^2+^ binds pp(dG)pp with an unexpected 2:1 stoichiometry according to our ITC experiment (SI Figure 3D). This interaction may be responsible for the different behaviors between ppGpp and pp(dG)pp hydrolysis at high Mn^2+^ concentration, i.e., pp(dG)pp hydrolysis is no longer faster with Mn^2+^ compared to Mg^2+^, and activity does not decline as [Mn^2+^] increase above 1.5 mM (compare Figures 5B and 4D). Because Mn^2+^ is not the physiological metal at the M2 site, mechanisms underlying these observations were not further investigated. Regardless, our data confirm that pp(dG)pp hydrolysis by SpoT*^Ab^* also requires Mg^2+^ or Mn^2+^ binding at a low-affinity site that presumably shares a similar environment to the M2 site for ppGpp hydrolysis (Figure 5B, top scheme).

**Figure 6.**
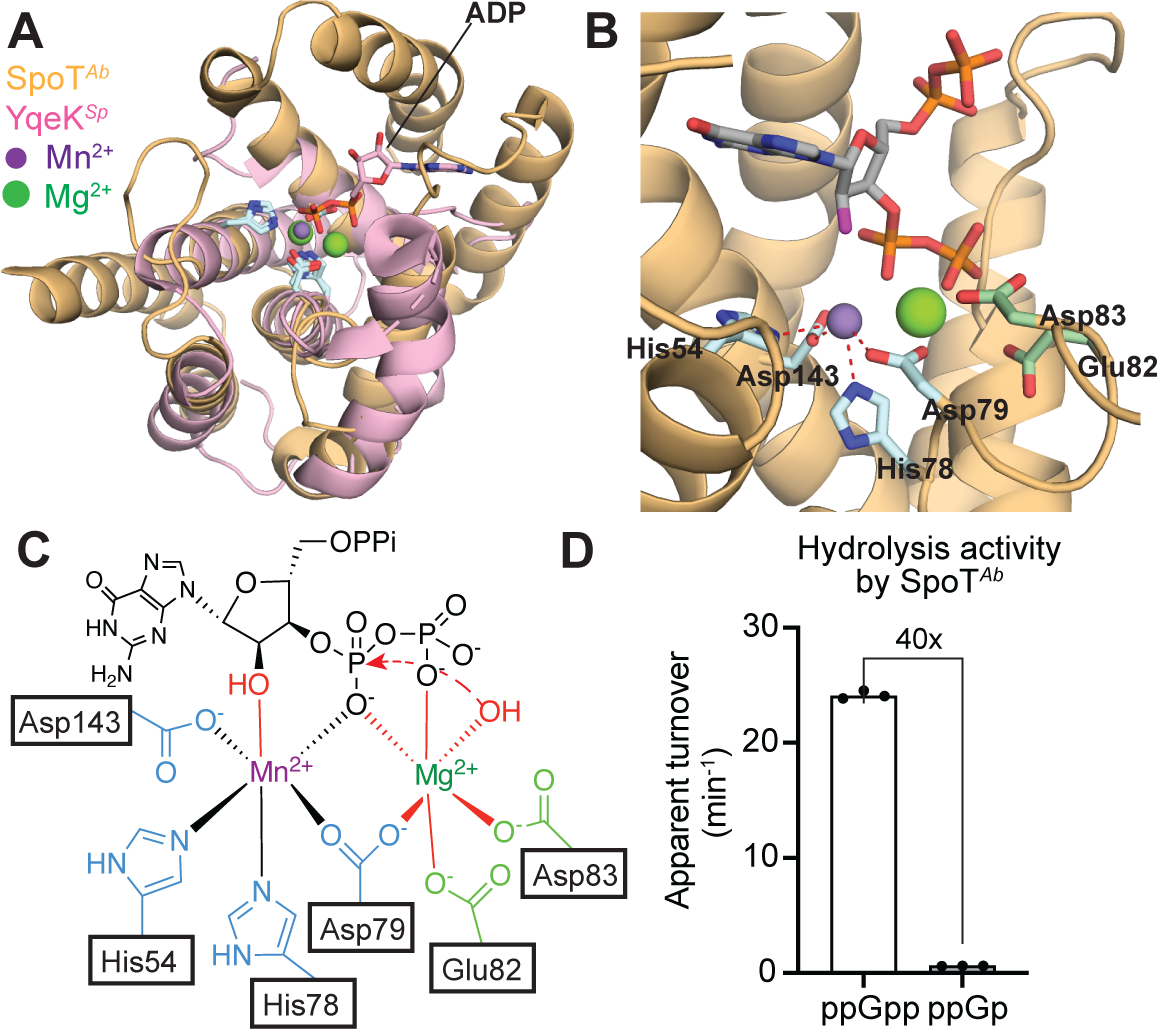
Proposed mechanism of ppGpp hydrolysis by SpoT^Ab^. (**A**) Structural alignment of SpoT*^Ab^* (orange, PDBID: 7QPR) and YqeK*^Sp^* (pink, PDBID: 8WMY). For clarity, YqeK*^Sp^* model includes only 5 closely aligned α-helices and loop/hinge regions in between. 2D2H motif of both proteins and ADP bound at YqeK*^Sp^* active center are shown in stick model; ppGpp at SpoT*^Ab^* active center is omitted. (**B**) Close-in view of SpoT*^Ab^*-ppGpp complex superimposed with the second Mg^2+^ in the YqeK*^Sp^* structure. (**C**) Proposed binuclear metallopyrophosphohydrolase mechanism of SpoT^Ab^. Interactions proposed based on the current study are shown in red bonds and arrows. (**D**) Hydrolysis kinetics of 500 μM ppGpp or ppGp by SpoT^Ab^ in the presence of 10mM MgCl_2_. To provide appropriate level of absorbance signal, ppGp hydrolysis was carried out with 1 μM SpoT^Ab^ compared to 100 nM for ppGpp.

We next varied the metal cation bound to M1-site in SpoT*^Ab^*. Among 4^th^ period divalent transition-metals, Cr^2+^ and Fe^2+^ are unstable to atmospheric oxygen at our reaction pH, while Cu^2+^ strongly inhibits the coupling of substrate hydrolysis and colorimetric readout by PK and lDH (data not shown). We hence included only Mn^2+^, Co^2+^, Ni^2+^ or Zn^2+^ in this assay. We added each cation at 100 μM, 1,000-fold molar excess relative to SpoT^Ab^, to drive its saturation of M1 site. We also used 1 mM EDTA to strip M1 site of any metal cation. In all five reaction conditions, 10 mM Mg^2+^ was provided to ensure saturated Mg^2+^ occupancy at the M2 site. Using ppGpp as the substrate, SpoT^Ab^ was inactive in buffers containing EDTA or Zn^2+^: the latter has been shown to potently inhibit Mesh1 by competing Mn^2+^ off the M1 site ^42^. Binding of Co^2+^ at M1 site doubled SpoT^Ab^ activity relative to the Mn^2+^-bound state (Figure 6C, compare blue to purple bar), while the neighboring element, Ni^2+^, provided less than 10% activity compared to Co^2+^ (Figure 5B, gray bar), suggesting that the coordination geometry of the M1 metal is critically important to the catalytic efficiency. Remarkably, despite much slower turnover, hydrolytic activity for pp(dG)pp holds a linear relationship with that for ppGpp across five buffer condition tested (Figure 6D and E). Thus, influence of the M1 metal on the activation energy is identical between ppGpp and pp(dG)pp hydrolysis. Our data strongly support the model in which ppGpp and pp(dG)pp adopt identical conformations for the rate-limiting step of their hydrolysis (Figure 5A, green arrows), and that slow *k_cat_* of pp(dG)pp hydrolysis dues to the decreased sampling of this productive conformation by the substrate. Taken together, our work revealed a potential reason why alarmone signaling rejects 2’-deoxynucleotide, namely, the 2’-OH group has a critical role in enabling alarmone degradation by RSH hydrolases.

## Discussion

In this study, we found that the SasB synthetase possesses an extremely low activity to synthesize pp(dG)pp from dGDP. SasB’s selectivity against 2dG nucleotides by 65,000-fold *k_cat_*/*K_m_* difference is stronger than that of RNA polymerase^34^, in consistence with our hypothesis that bacteria avoid synthesizing 2dG alarmones because of a strong negative-selection pressure. With the help of CCMS, we found that SpoT, and perhaps all RSH hydrolases, are deficient in hydrolyzing 2’-deoxy 3’-pyrophosphorylated substrates. This finding opens the possibility that the inability to degrade (p)pp(dG)pp constrains alarmone signaling from utilizing 2dG nucleotides. To delineate the role of 2’-OH in the hydrolase activity of SpoT, we interrogated the divalent metal-cation dependence of both ppGpp and pp(dG)pp hydrolysis and showed that SpoT is a binuclear metallophosphohydrolase. Based on our new results, we present an updated model for the catalytic mechanism of RSH hydrolases and an outlook regarding the existence and function of (p)pp(dG)pp in biology.

### New insights to RSH hydrolase mechanisms

The ability to degrade alarmones is arguably more important than the ability to synthesize these molecules. As a widespread feature across the bacterial kingdom, an organism often encodes more than one synthetases, each specialized for producing alarmones under certain starvation conditions, but relies on a single hydrolase for alarmone degradation. Deletion of the lone hydrolase in such organisms lead to loss of viability in many species, making these hydrolase domains potential drug targets. In the past two decades, crystallographic studies have provided high-resolution snapshots for many single-domain SAHs or bifunctional, long-RSH proteins, some with ppGpp, the substrate, present at the hydrolase active center. Despite this progress, the precise catalytic mechanism of alarmone hydrolysis has remained elusive. Specifically, it is unclear whether the hydrolysis occurs by water directly attacking the 3’-α-phosphate (direct mechanism) or a protein nucleophile attacking the 3’-α-phosphate followed by subsequent hydrolysis of a pyrophosphorylated protein intermediate (ping-pong mechanism). Additionally, all RSH hydrolases require an invariant Glu-Asp (ED) diad for hydrolase activity ^8^, whose side-chain carboxylates are about 6Å away from the 3’-α-phosphorus or the active-center (M1-site) Mn^2+^ (Figure 6A, highlighted in green)^34^. Little is known about the catalytic role of this ED diad.

In this study, we show that SpoT*^Ab^*, and perhaps other members of the RSH hydrolase family are binuclear metallopyrophosphohydrolases requiring a Mg^2+^ with millimolar EC_50_ in addition to the 2H2D-chelated Mn^2+^ for catalysis. Importantly, all RSH hydrolases belongs to a clade of HD-domain superfamily featuring the 2H2D metal-binding site^43^. A non-RSH member of this clade, *Streptococcus pyogenes* YqeK, a hydrolase of diadenosine tetraphosphate (Ap4A), has recently been crystallized with two Mg^2+^ around the catalytic center (PDBID: 8WMY, Figure 6A, pink)^44^. While the first Mg^2+^ binds at the 2H2D site, the second Mg^2+^ is sandwiched between ADP and an Asp residue from the 2H2D motif (Figure 6A). All *YqeK^Sp^* α-helices embedding the 2H2D motif superimposed well to those in SpoT*^Ab^* with an RMSD = 3.0 Å across 80 Cα atoms (Figure 6A, orange, see Figure 3A for sequence alignment). Intriguingly, superimposing the second Mg^2+^ in YqeK*^Sp^* with SpoT^Ab^-ppGpp structure (Figure 6B, green sphere) revealed a metal binding site that would be readily coordinated by Asp79, both 3’-phophates of ppGpp, and likely the ED diad consisting of Glu82 and Asp83 (Figure 6B). We therefore propose that this Mg^2+^ approximates the M2 site of SpoT*^Ab^* and that the ED diad’s essential function is to accommodate the M2-site divalent cation (Figure 6B and 6C). In support of the role of 3’-phosphate as the M2-site ligand, we further confirmed that SpoT^Ab^ is unable to hydrolyze a substrate missing the 3’-β-phosphate, ppGp (**8**) (Figure 6D). Our study also revealed a critical role of the 2’-OH group in stabilizing the catalytically-productive conformation of SpoT*^Ab^*-ppGpp complex. This effect likely comes from direct coordination of 2’-OH to the M1-site Mn^2+^ as was indicated by the short O-Mn distance (3.5-4.0 Å) in the crystal structure (Figure 6B and C). Since the M1-site Mn^2+^ would reach a valency of six upon coordination by 2’-OH, the nucleophile attacking the 3’-α-phosphate should be activated by the M2-site Mg^2+^. Because 3’-α-phosphate is apparently not accessible by any protein nucleophile from the correct geometry, we speculate this nucleophile should be a hydroxide deprotonated from a water molecule (Figure 6C). Notably, in substrate-free RSH-hydrolase structures, the loop bearing the ED diad retracts away from the active center, suggesting flexibility between M1 and M2 sites unless bridged by the ppGpp substrate ^38^. Without 2’-OH latching on the Mn^2+^, geometry between 3’-α-phosphate and the attacking hydroxide may deviate from the optimum, thus resulting in a drastic decrease of *k_cat_* (Figure 6A, red arrow). Ultimately, testing of our proposed mechanism would require a high-resolution structure of a RSH hydrolase bound to a transition-state mimic ^45^.

### Potential existence and roles of pp(dG)pp and its synthetase

Our study reveals the potential lack of competent pp(dG)pp degradation machinery in bacteria, which should contribute, at least in part, to the negative selection against 2dG alarmones. It is also noteworthy, however, that although SpoT*^Ab^* hydrolyzes ppGpp with *k_cat_*/*K_m_* 1,000-fold faster than pp(dG)pp, it is not nearly as selective compared to SasB. The more stringent selection on the synthetase side of alarmone regulation suggests additional selection pressure driving this specificity. Upon acute starvation, both *E. coli* and *B. subtilis* produce (p)ppGpp at millimolar levels^19, 35^. In *B. subtilis*, this high level of alarmone production is achieved by the strong synthetase activity of SasB upon activation by pppGpp ^28^. Consumption of GTP and GDP by this process, as well as the inhibition of guanylate kinase by ppGpp^46^, leads to depletion of guanosine-5’-nucleotides. Low GTP level drives transcription reprogramming that adapts the bacterium to the low-nutrient condition^36^. Nevertheless, had SasB synthetases not specifically exclude dGTP or dGDP, its activation would also deplete dGTP and potentially interrupt DNA replication. This effect is reminiscent of thymine-less death and may have severe fitness consequences ^37^. Thus, we speculate that feed-forward synthetases like SasB undergo the strongest selection pressure to avoid dGDP to insulate mass production of alarmones from 2’-deoxyribonucleotides and DNA replication. Testing this speculation would entail data regarding GDP vs dGDP specificity of a housekeeping synthetase (e.g., SpoT*^EC^*), although we have not been able to detect pp(dG)pp synthesis by any RSH synthetase other than SasB.

To our best knowledge, no RSH synthetases have previously been shown to utilize 2’-deoxyribonucleotide as diphosphate acceptor. Nevertheless, recent studies have revealed non-canonical pyrophosphokinase activities for RSH synthetases in toxin-antitoxin modules. Toxicity of these proteins are achieved by rapid modification of adenosine nucleotides or tRNA to disrupt energy production or translation. Under physiological conditions, these toxins are activated by either secretion to another bacterium cell to antagonize the growth of the latter, or by recognition of phage proteins to halt phage replication and protect the host bacterial population ^21, 24^. Since a strong, specific (p)pp(dG)pp synthetase activity would almost certainly deplete dGTP and disrupt DNA synthesis, we speculate that if such an enzyme exists, it most likely to be encoded in a toxin-antitoxin (TA) module to carry out similar phage-defense or interspecific antagonism functions. Importantly, single mutation is known to alter specificity of an enzyme to a structurally similar substrate. For example, the R132H mutant of human cytosolic isocitrate dehydrogenase I (IDH1) adopts a specific 2-hydroxyglutarate dehydrogenase activity associated to glioma and glioblastomas^47^. Thus, a stronger and more specific (p)pp(dG)pp synthetase is likely accessible from canonical RSH synthetases via a relatively short evolutionary path. Once discovered, these enzymes may serve as powerful tools to, for example, perturb 2’-deoxyribonucleotide metabolisms for further investigation of the mechanisms of thymine-less death.

## Acknowledgement

This work was supported by the Pathway of Independent Award from NIH (R00GM135536) and a Research Grant from Robert Welch Foundation (I-2171) to B.W.. B.W. is a Southwestern Medical Foundation Scholar in Biomedical Research. We thank Dr. Andrew Lemoff at the UTSW Proteomics Core and Dr. Feng Lin as the UTSW Nuclear Magnetic Resonance Core for carrying mass spectrometry and NMR analysis, respectively. We thank Dr. Elliott M. Ross and Dr. Kimberly A. Reynolds at UT Southwestern Medical Center for helpful discussions.

## Author contributions

B.W. conceived the project and designed the research. R.W.Z., I.J.G and B.W. performed proteomic and biochemical experiments and analyzed the data. Y.H. carried out NMR analysis of new compounds and analyzed the data. R.W.Z. and B.W. wrote the manuscript with input from all authors.

---

**SI Figure 1.**
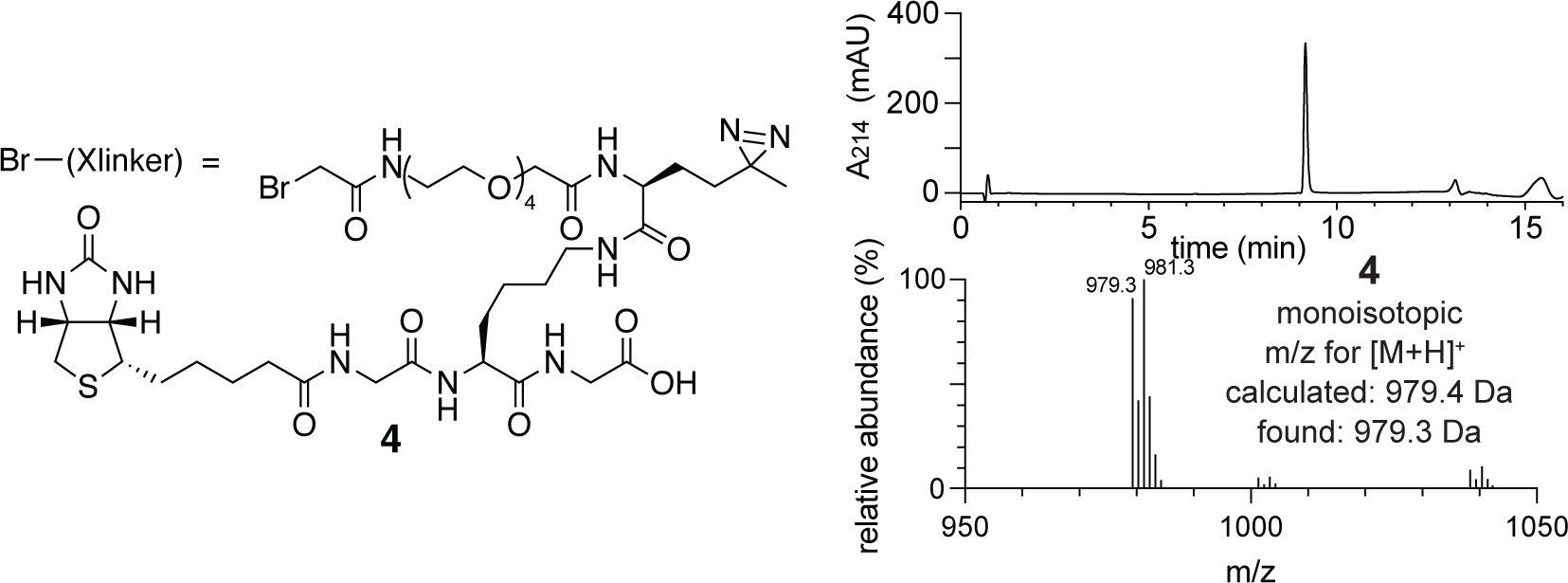
Characterization of bromoacetylated crosslinking peptide (**4**, panel **A**). C18 HPLC chromatogram (right top) ESI-MS analysis in positive mode (right bottem) are shown.

**SI Figure 2.**
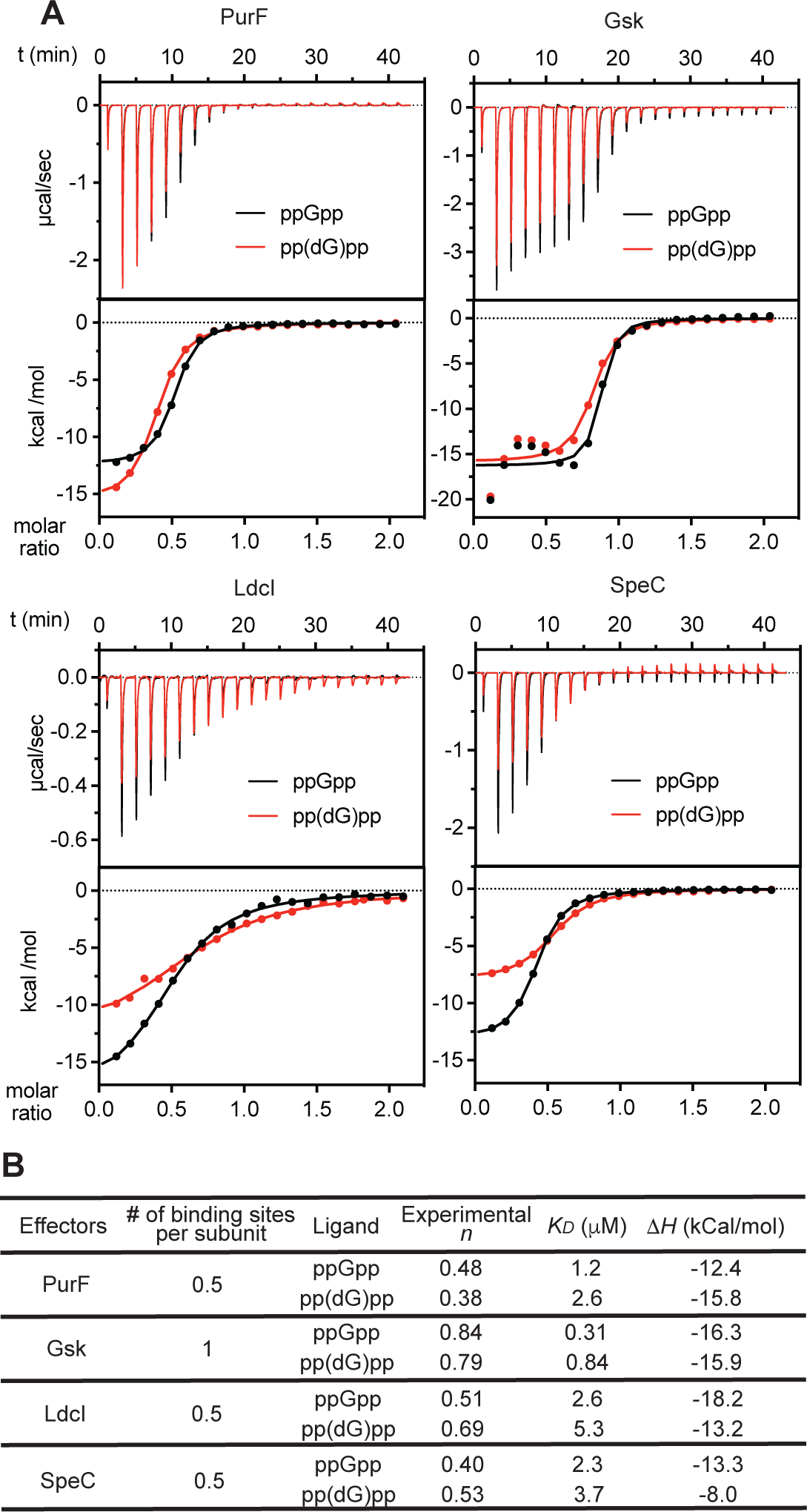
pp(dG)pp retains interaction with known ppGpp effectors. (**A**) ITC traces (top) and fitting isotherms (bottom) for indicated protein with ppGpp (black) or pp(dG)pp (red). 100 μM of PurF, Gsk and SpeC and 21.6μM of LdcI (counted as monomers) were titrated with 10x concentration of ligand. (**B**) fitting parameters for datasets in (**A**).

**SI Figure 3.**
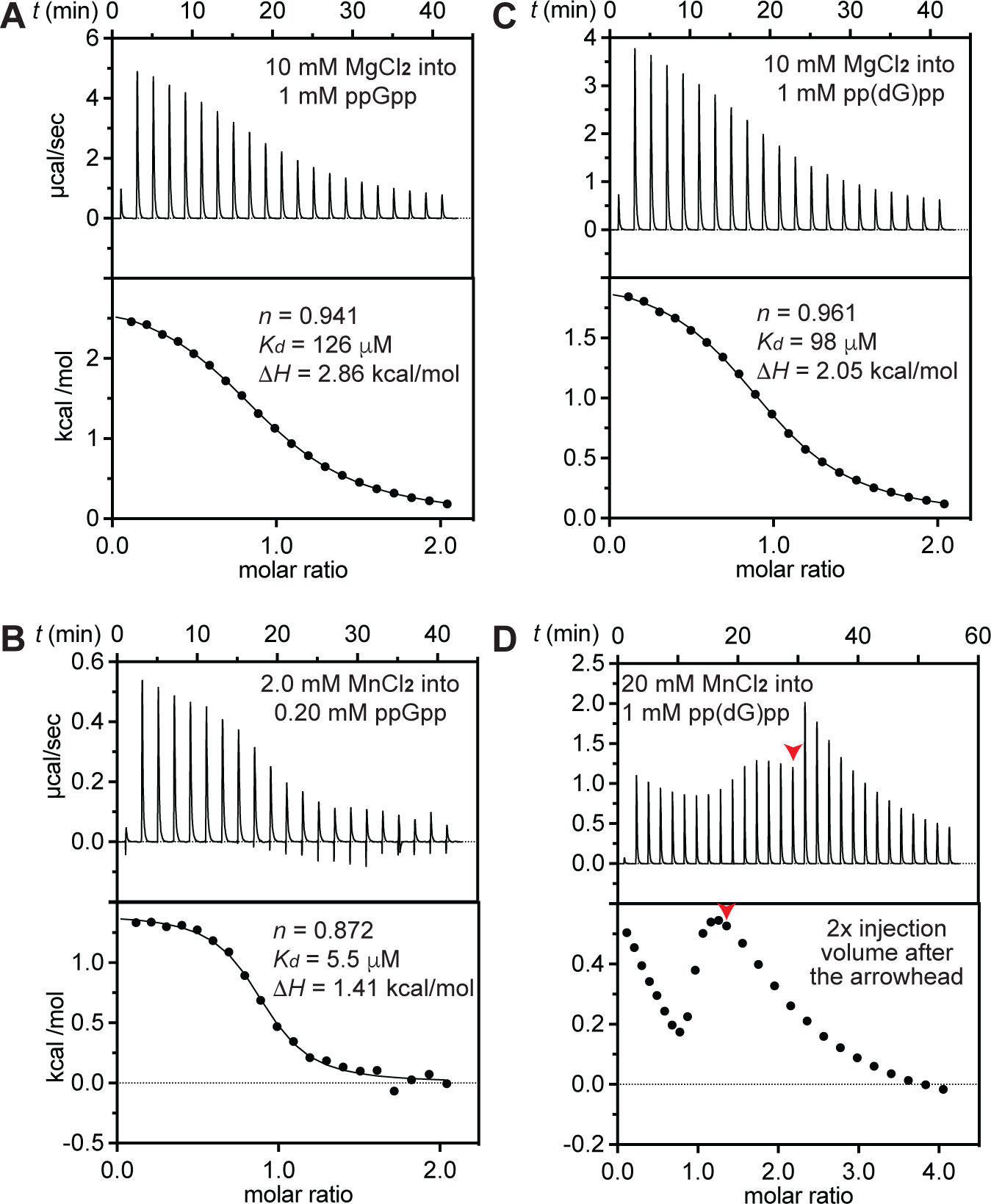
ITC of ppGpp and pp(dG)pp using indicated divalent metal salts. Fitting results are shown for panels **A**-**C**. Results in panel **D** indicates that MnCl_2_ binds pp(dG)pp at 2:1 as the heat signal declines towards to the baseline after a molar ratio of 2.0, but we cannot deconvolute the *K_D_* of each binding event.

## Methods

### Contact for Reagent and Resource Sharing

Questions about or requests for methods, strains, and resources generated in this study can be directed to Boyuan Wang.

### Media and Culture Conditions

*Escherichia coli* was grown in LB (10 g/L NaCl, 10 g/L tryptone, 5 g/L yeast extract). Agar plates were incubated at 37 °C unless noted otherwise. Large cultures (> 10 mL) were grown in baffled Erlenmeyer flasks at 37 °C and 200 rpm on an orbital shaker, and OD_600_ measurements were taken using a 1-cm plastic cuvette. If necessary, the culture was diluted to OD_600_ < 0.4 and the dilution was factored in. Appropriate antibiotics was included at following concentrations for plasmid maintenance: Carbenicillin—100 mg/mL; kanamycin—50 mg/mL; chloramphenicol—25 mg/mL.

### Plasmid construction

All primers and plasmids used are listed in Table S1 and S2, respectively

*Sub-cloning*: We used Gibson-assembly to ligate vector and insert DNA each of which were PCR-amplified using primers containing appropriate overhangs to ensure ligation specificity. All PCR reactions were performed using the Q5 High-Fidelity DNA Polymerase (NEB) kit following the instructions enclosed. PCR products were purified with agarose gel-electrophoresis followed by gel extraction prior to ligation (Zymoclean Gel DNA Recovery Kit). Circular plasmids were extracted from positive transformants and the presence of correct insert confirmed through Sanger sequencing.

The *A. baumannii* SpoT open reading frame (ORF) was sub-cloned into pET SUMO vector (Invitrogen) immediately downstream of the His_6_-SUMO ORF. The *E. coli* ldcI ORF was sub-cloned into pCfa vector upstream of the C-Cfa intein-His_6_ ORF^1^.

### Recombinant protein production and purification

*Protein expression*: all recombinant proteins were expressed in *E. coli* BL21(DE3) harboring the corresponding expression plasmid. Expression strains were grown in LB containing the appropriate antibiotics at 37 oC to OD_600_ = 0.6. The culture was then cooled to 18 oC and induced by the addition of 400 µM IPTG. Cells were harvested 16 to 24 hours post-induction. ldcI was underwent fast induction with 400 µM IPTG, at 30oC and harvested 3 hours post-induction.

*Purification of His-tagged proteins*: His-tagged proteins were purified following the standard protocol described below unless otherwise noted. SasB was purified by substituting 150mM NaCl with 300 mM KCl in all buffers. SpoT^Ab^ was purified by substituting 150mM NaCl with 300mM NaCl and 150 mM KCl. The cell pellet from 1L of expression culture was resuspended in 25 mL lysis buffer containing 50 mM Tris-HCl pH 8, 150 mM NaCl, 1 mM TCEP, 20 µg/mL lysozyme, 10 units Benzonase (MilliporeSigma) and 1 mM PMSF. Cells were disrupted through sonication and the lysate was cleared at 12,000 *g* for 1 hour. Cleared lysate was applied to 3 mL Ni-NTA resin equilibrated with the lysis buffer and allowed to flow through by gravity. The Ni-NTA resin was washed with 5x column volumes of wash buffer 1 (50 mM Tris-HCl pH 8, 150 mM NaCl, 10 mM imidazole and 1 mM TCEP) and 10x column volumes of wash buffer 2 (50 mM Tris-HCl pH 8.0, 150 mM KCl, 25 mM imidazole 1 mM TCEP). Bound protein was eluted with 3X column volumes of elution buffer (50 mM Tris-HCl pH 8.0, 150 mM NaCl, 400 mM imidazole and 1 mM TCEP. For SpoT^Aba^, 50mM EDTA was used instead of imidazole. Imidazole/EDTA was dialyzed out using Amicon Ultra-15 Centrifugal Filters with the appropriate MW cutoff by repetitively concentrating down the eluate and re-dilute with Imidazole/EDTA-free buffer.

*Removal of affinity tags*: All proteins except SasB were removed from affinity tags prior to usage for biochemical/biophysical experiments. To remove the N-terminal His_6_ tag of recombinant proteins expressed from the pET28b vector, thrombin (Sigma) was reconstituted in 20 mM Tris-HCl pH 8.0, 150 mM NaCl and 20% (v/v) glycerol at 100 U/mL. Eluates off the Ni-NTA column were dialyzed against 20 mM Tris-HCl pH 8.0 and 150 mM NaCl, and then treated with thrombin (10 U per µmol cleavage sites) at RT overnight. Upon confirming completion of cleavage using SDS-PAGE, the protein was concentrated, treated with 10 mM TCEP, and refined over a Superdex-200 increase (10/300) column. To remove the N-terminal His_6_-Sumo tag, proteins off the Ni-NTA column were exchanged into 20 mM Tris-HCl pH 8.0, 150 mM NaCl and 1 mM TCEP. The tagged protein was then treated with Sumo protease (10 µg per µmol cleavage sites) at RT for 2 hrs. The cleavage mixture was subjected to a reverse Ni-NTA process to clean up residual uncleaved protein. Flow-through containing cleaved protein was then concentrated and refined over a Superdex-200 increase (10/300) column.

*Size-exclusion chromatography (SEC)*: A Superdex-200 Increase 10/300 column (Cytiva) was used. SEC of all recombinant proteins with the exception of SasB was run in HBST buffer containing 20 mM HEPES-Na pH 7.5, 150 mM NaCl and 1 mM TCEP. Up to 10 mg protein was injected in 1 mL and peak fractions were combined. SasB was run in 50mM Tris-HCl pH 8, 300mM KCl and 1mM TCEP. Complete purification was confirmed with SDS-PAGE.

### Synthesis and purification of nucleotides bearing 3’-diphosphate or 3’-phosphate

Purified *B. subtilis* SasB was used to synthesize pp(dG)pp **1**, ppGpp-3βS **2**, pp(dG)pp-3βS **3**, and ppGpp. The reactions were conducted in 2mL containing 20mM Tris buffer(pH 9.1 or 8), 150 mM NaCl, 25mM MgCl2 and 1mM TCEP. Final pH was Tris-base 9.1 for 2’-deoxy and dG4P3γS and Tris-HCl 8.0 for all 2’-OH guanine nucleotide synthesis. 250µM pppGpp was included to activate SasB ^2^. Each reaction included 20 mmol (10 mM, 1.0 eq) appropriate guanine nucleotide and 24 mmol (1.2 eq) ATP or ATPγS. Final concentrations of SasB were 50 μM for pp(dG)pp and 100 μM for dG4P3γS synthesis, which reached completion in 72 hours, and 1 μM for 2’-OH guanine nucleotide synthesis, which reached completion in 3 hours. To monitor reaction progress, 1 mL of reaction was analyzed on MonoQ 5/50 10-40%B in 15 mL. All MonoQ runs were complete with Buffers A: 5 mM Tris-HCl 8.0 and Buffer B: 5mM Tris-HCl 8.0 and 1mM NaCl. Upon completion, the reaction was vortexed with 300uL chloroform for 1 minute, and spun at 21,000G for 5 min. The aqueous layer was diluted 20-fold in water and resolved on MonoQ 10/100 10-40%B in 80mL. Fractions containing the product were combined, lyophilized and re-dissolved in water to 5mL. LiCl powder was then added to 1 M concentration, and the products were precipitated by addition of 4x volume of ice-cold 100% ethanol. After staying in ice-water bath for 30 min, precipitate was spun down at 3,300G for 5 min and supernatant was aspirated. The nucleotides were washed with 1x volume of 95% EtOH. Sonication was used to disperse the precipitate for better removal of the leftover LiCl. The product was spun down and washed 3x before being re-dissolved in water and quantified based on light absorption at 252 nm and then lyophilized. Dried, colorless powder was dissolved with Milli-Q water into 100mM stocks and stored frozen -80°C.

ppApp was synthesized using *P. aeruginosa* Tas1(230-439), also known as Tas1^Tox^, as previously described ^3^. Briefly, 45μmol ADP was added in 9 portions to 2.5mL solution of 50 μmol ADP and 50 pmol Tas1^tox^ in 5 mL 20 mM HEPES 7.4, 150 mM NaCl, 20 mM MgCl_2_ and 1 mM TCEP-Na at 37°C with vigorous stirring, we added 45 µmol ATP in 9 portions over 10 minutes. After another 5 minutes of incubation at 37°C, the reaction was complete and Tas1tox was inactivated with 2 mL chloroform. The aqueous phase was isolated, diluted to 25 mL with water, and ppApp purified similarly using MonoQ 10/100, precipitated with LiCl and EtOH as described above.

pp(dA)pp **7** was synthesized following strategy developed by Haas, et.al ^4^. 2’-deoxyadenosine (2dA) monohydrate (211 mg, 784 μmol) was coevaporated using dry MeCN (4 x 15 ml). Afterwards diphosphoryl chloride (1.25 mL, 2.28 g, 9.0 mmol, 11.5 eq) was added in one portion at −35°C to dissolve all 2dA. The reaction was immediately protected with dry N_2_ and stirred for 3 h at 0°C. Afterwards, the reaction was suspended in 20mL dry ether, vortexed and sonicated at 0°C. Dark gray insoluble content was collected by centrifugation, chilled in ice water, and ether decanted. The insoluble was then dissolved by addition of ice-cold 0.5M NaHCO_3_ solution in 3-mL portions. A total 30 mL solution is needed to bring pH above 7.0. The resulted solution was degassed, filtered, and 2’-deoxyadenosine-3’-5’-bisphosphate, p(dA)p, was purified using MonoQ 10/100 and precipitated with LiCl and EtOH as described above. This crude p(dA)p (136 μmol, 17%) was dissolved in 5 mL water and exchanged into tetrabutylammonium (TBA) salt. To do so, we pre-treated a Hi-Trap CM Sepharose FF column (10 mL) with 2 mL 1 M TBA bromide solution follow by washing with 10 mL water. Flowing the p(dA)p solution through the column gave rise to a solution of p(dA)p•TBA_3_, which was lyophilized and re-dissolved in dry DMF. Note that the compound changed from colorless to brown following lyophilization but this did not affect the subsequent reaction. To a 1mL DMF solution containing 40 μmol p(dA)p•TBA_3_ and 25.4 mg 4,5-dicyanoimidazole (215 μmol, 5.4 eq) was added 600 μL 0.2 M dry-DMF solution of (bis((9H-fluoren-9-yl)methyl) diisopropylphosphoramidite (iPr_2_NP(OFm)_2_, 120 μmol, 3.0 eq). After stirring the solution at RT for 15min, the reaction was treated with 25.6 mg 3-chloroperbenzoic acid (*m*CPBA, 77%, 191 μmol, 4.8 eq, pre-dissolved in 500 μL dry DMF) at 0°C for 15 min followed by 1,8-diazabicyclo(5.4.0)undec-7-ene (250 μL, large excess, pre-dissolved in 2 mL dry DMF) at RT for 15 min. Crude product was precipitated with 30 mL cold ether, collected by centrifugation and washed with 10 mL cold ether. The precipitate was dissolved in 2 mL 50 mM Tris-HCl 8.0, filtered and the final product, 2’-deoxyadenosine-3’-5’-bis(diphosphate), pp(dA)pp, was purified and using MonoQ 10/100 and precipitated with LiCl and EtOH as described above. Isolated yield was 8.2 μmol, 20.5% for the second step and 3.4% overall.

ppGp **8** was obtained by cleaving the 3’-β phosphate of ppGpp with RNase T2. The reaction occurred in 2 mL of 20mM sodium acetate buffer pH 4.6, 150 mM NaCl, and 1mM TCEP. 20 μmol ppGpp was cleaved with 1U of T2 RNAse and monitored on MonoQ/5 50 until completion. Chloroform precipitation of protein and aqueous extraction was performed as above. Sample was similarly applied to MonoQ 10/100 and eluted. LiCl and EtOH precipitation, quantification, and lyophilization was repeated as above.

### Solid-phase peptide synthesis

Note that all equivalences are relative to the loading capacity of the solid support resin.

The general procedure for a coupling-deprotection cycle: 10 eq. Fmoc protected amino acid and 9.5 eq. of 1-[Bis(dimethylamino)methylene]-1H-1,2,3-triazolo[4,5-b] pyridinium 3-oxide hexafluorophosphate (HATU) dissolved in N,N-dimethylformamide (DMF) to achieve a final concentration of 0.4M amino acid and 0.38M HATU. 20 eq. Of diisopropylethylamine (DIEA) was then added and the mixture was applied to the solid support and left to react at room temperature (RT) for 20 minutes, followed by 3 washes with DMF, 2x 8 min deprotection with 20% (v/v) piperidine in DMF and 5 washes with DMF. Biotin was coupled following the same procedure but without deprotection. For coupling of Fmoc-photoMet, usage of the amino acid, HATU and DIEA were reduced to 2.0, 1.9, and 10 eq., respectively, and coupling time extended to 1 hour at RT.

The Biotinyl-GKG structure first synthesized at 0.1-mmol scale on Fmoc-Gly-Wang resin (Sigma-Alrich). The resin was swelled in DMF prior to the first coupling. Coupling-deprotection cycles were performed with Fmoc-Gly-OH, Fmoc-Lys(Alloc)-OH, and Fmoc-Gly-OH (in this order), and the N-terminal amine was biotinylated. Thereafter, The Alloc protecting group was quantitatively removed by a 20-minute, RT treatment with 598 µL of phenylsilane (4.85 mmol) and 45.63 mg (39.5 µmol) of tetrakis(triphenylphosphine)palladium(0) in 4 mL of DCM. The resin was then thoroughly washed with dichloromethane (DCM), air dried, and stored in a desiccator for further use.

Syntheses was continued on a 0.01-mmol scale. Resin bearing the Biotinyl-GKG fragment was weighed into individual vessels and re-swelled with DMF. Fmoc-photoMet-OH were coupled to the lysine side chain followed by the linker residue: Fmoc-NH-PEG4-OH (ChemPep, Cat# 280117). The N-terminus of the linker residue was then bromoacetylated by adding an 1 mL DMF solution containing 40 mL bromoacetic anhydride and 50 mL DIEA (30 eq) for a 10-minute incubation at RT. Finally, the resin bearing each peptide was washed thoroughly with DCM, air dried, and the product peptide was cleaved from the resin with 3 mL neat trifluoroacetic acid (TFA) containing 2.5% (v/v) water and 2.5% (v/v) triisoproprylsilane at 60 °C for 8 min. TFA in the resulting solution was evaporated by a stream of nitrogen gas and the residual was triturated in cold diethyl ether to precipitate the peptide. The crude peptide was pelleted, washed once with diethyl ether and re-dissolved in 50% (v/v) acetonitrile in water containing 0.1% TFA and lyophilized. The crude peptide was refined over a semi-preparative reverse phase (RP)-HPLC (Discovery BIO wide pore C18 column: 10 x 250 mm, 5 µm) using a linear gradient of solvent A (0.1% TFA in 90% acetonitrile) and B (0.1% TFA) with the percentage of B increasing from 5% to 25% over 10 minutes. Flow rate was 6 mL/min. Pure fractions of the product peptide were combined and lyophilized.

### Capture compound synthesis

Capture compounds were synthesized by conjugating 1.2 equivalence bromoaceylated peptide to 2 μmol ppGpp-3βS at pH = 6.2 or pp(dG)pp-3βS and at pH = 7.5. Reactions were carried out at 42 °C for 1 hr and then treated 5 μmol sodium 2-mercaptoethanesulfonate (MESNa, Sigma-Aldrich) for 15 minutes. Product was purified using a MonoQ (5/50) column at RT °C using a linear gradient of buffer A (5 mM Tris-HCl pH 8.0) and buffer B (5 mM Tris-HCl pH 8.0, 1M NaCl), with the percentage of buffer A increasing from 20% to 40% in 10mL.

### Isothermal titration calorimetry (ITC)

All ITC experiments were performed in a MicroCal PEAQ-ITC (Malvern) instrument thermo-equilibrated at 20°C with pure water in the reference cell. All ligands and proteins were suspended in the same buffer containing 20 mM HEPES-Na pH 7.5, 150mM NaCl, 2mM TCEP. The cell(protein) volume was 200uL. Ligand and protein concentrations are given in S Table #. The cell(protein) volume was 200uL. For all titrations, the ligand was injected at 1.9 µL/injection for altogether 21 injections. ITC data were processed using MicroCal PEAQ-ITC Analysis Software (Malvern), which automatically integrates heat signal and calculates the molar enthalpy change (DH_m_) and the overall ligand-to-receptor molar ratio ([L]/[R]) at each injection. Titration trace was corrected by using the default subtract baseline function. ΔH_m_ was corrected by using the subtract offset function. All titrations were fitted against a sigmoidal Δ *H*_m_ to molar ratio relationship in a single-site model with both ligand-to-protein stoichiometry (*n*) and *K*_D_ as variables.

### Effector capture and enrichment from *E. coli* lysate

*E. coli* MG1655 was cultured at 37°C in 200-mL batchs in 1-L baffled flask with vigorous shaking to OD_600_ = 0.40. M9-Glucose SILAC medium containing 12.8 g/L Na_2_HPO_4_•7H_2_O, 3.0 g/L KH_2_PO_4_, 0.50 g/L NaCl, 0.4% (w/v) glucose, 1 mM MgCl_2_, 30 μM CaCl_2_, 20 μM FeSO_4_-EDTA and 0.50 g/L regular NH_4_Cl (light medium) or 0.51 g/L NH_4_Cl (15N, 99%, Cambridge Isotope Laboratories, heavy medium) were used. Lysates were prepared in 20 mM HEPES-Na 7.0, 300 mM NaCl and 1 mM TCEP at approximately 10 mg / mL protein. Then, for each capture compound (**2** or **3**), two reactions were assembled in adjacent wells on a 96-well plate chilled on ice. The effector-capture reaction consisted of 800 μg protein from the light lysate, 100 μM capture compound, 1.8 mM MgCl_2_, and 0.2 mM MnCl_2_. The control reaction consisted of 800 μg protein from the heavy lysate, 100 μM capture compound, 5 mM ppGpp, 6.3 mM MgCl_2_ and 0.7 mM MnCl_2_. After exposure to 365-nm UV lamp for 2 minutes, reactions were then combined and extensively exchanged in a 10kDa-MWCO concentrator into a buffer containing 20 mM HEPES-Na pH 7.5 and 200 mM NaCl. The final concentrate (approximately 100 μL) was diluted with 400 μL RIPA buffer (50 mM Tris 7.5, 150 mM NaCl, 0.1% SDS, 0.5% sodium deoxycholate, 1% Triton X-100) and incubated overnight with 200 μL MyOne Streptavidin C1 dynabeads (Thermo Fisher Scientific) at 4 °C. On the second day, the protein in supernatant (flowthrough) was sampled for Western blotting and then removed, and the dynabeads were washed at 4 °C twice with RIPA buffer, once with 0.1 M Na_2_CO_3_, once with 100 mM Tris 8.0, 4 M guanidinium chloride (GuHCl) and twice more with RIPA buffer. Bound protein was eluted through boiling the dynabeads in 40 μL SDS-PAGE loading dye containing 2 mM biotin. The eluate was lyophilized and re-diluted using water to 15μL, and resolved on 12% TGX-PAGE (Bio-Rad, Mini Protean) at 100V for 1 hour.

### LC-MS2 based proteomics

The PAGE gel containing proteins eluted from streptavidin pulldown was stained with Coomassie Brilliant Blue (CBB). The gel lane for each eluate was excised and regions below the 15-kDa or above the 200-kDa marker were trimmed off. The gel lane was diced into 1-mm pieces, and then treated with 10 mM DTT at 56 °C for 1 hr followed by 55 mM iodoacetamide at RT for 1 hr in the dark. After a brief wash with 0.1 M NH_4_HCO_3_, gel pieces were dehydrated with acetonitrile, dried over a lyophilizer, and re-hydrated with a minimal volume of 6 ng/μL trypsin in 0.1 M NH_4_HCO_3_ for digestion at RT overnight. Thereafter, tryptic fragments were extracted by four shrinking-swelling iterations. In each iteration, gel pieces were first dehydrated with 5% formic acid in 50% (v/v) acetonitrile (first two iterations) or neat acetonitrile (second two). Supernatant containing tryptic fragments was then removed for collection and gel pieces were re-swelled with 0.1 M NH_4_HCO_3_ in water. The combined supernatant was dried in a speedvac and reconstituted in 10 μL 0.1% formic acid. 5 μL was used for LC-MS2 analysis.

Tryptic fragments were injected onto an Q Exactive HF mass spectrometer coupled to an Ultimate 3000 RSLC-Nano liquid chromatography system. Samples were injected onto a 75 um i.d., 15-cm long EasySpray column (Thermo) and eluted with a gradient from 0-28% buffer B over 90 min. Buffer A contained 2% (v/v) ACN and 0.1% formic acid in water, and buffer B contained 80% (v/v) ACN, 10% (v/v) trifluoroethanol, and 0.1% formic acid in water. The mass spectrometer operated in positive ion mode with a source voltage of 2.5 kV and an ion transfer tube temperature of 300 °C. MS scans were acquired at 120,000 resolution in the Orbitrap and up to 20 MS/MS spectra were obtained in the ion trap for each full spectrum acquired using higher-energy collisional dissociation (HCD) for ions with charges 2-8. Dynamic exclusion was set for 20 s after an ion was selected for fragmentation.

Raw MS data files were analyzed using Proteome Discoverer v3.0 SP1 (Thermo), with peptide identification performed using a tryptic search with Sequest HT against the *E. coli* reviewed protein database from UniProt. Fragment and precursor tolerances of 10 ppm and 0.02 Da were specified, and three missed cleavages were allowed. Carbamidomethylation of Cys was set as a fixed modification, with oxidation of Met set as a variable modification. Quantification was provided for peptides with only ^14^N-containing amino acids (light) and those with only ^15^N-containing amino acids (heavy). The false-discovery rate (FDR) cutoff was 1% and XCorr-score cutoff was 1.8 for all peptides.

### LC-MS2 data processing

With each peptide group, Sequest returns quantifications of the light and heavy forms. A few peptides were returned with neither form quantified which were discarded. L/H ratio was directly calculated for peptides with both light and heavy forms quantified. We then sorted all peptides by their L/H ratio. To avoid averaging bias introduced by extreme L/H values, we arbitrarily reassigned the L/H values of all the top 2% peptides and all peptides with only light form quantified with the L/H value of the 98^th^ percentile peptide. We similarly reassigned the L/H values of the bottom 2% peptides with the L/H value of the 2^nd^ percentile peptide. We then calculated log_2_(L/H) for all peptide groups by averaging log_2_(L/H) values of all peptide groups (equal weight) belonging to that protein. In addition to log_2_(L/H), we archived the total number of PSMs (#Totpep), peptide groups (#Unipep) and peptide groups of arbitrarily assigned L/H values (#Arbpep). Only proteins reliably detected (criteria: #Unipep >= 3 OR (#Unipep = 2 AND #Arbpep = 0)) in BOTH replicates were considered for hit selection and are shown in the log_2_(L/H) correlation plot. For each capture compound, all reliably detected proteins were sorted by log_2_(L/H) averaged from two replicate datasets and the top 20% were arbitrarily chosen as ppGpp-binding hits. These log_2_(L/H) values were also used for the correlation plot between ppGpp and pp(dG)pp capture compounds (Figure 2E). In contrast to the log_2_(L/H) correlation plot, all proteins with as few as one PSM detected in either replicate are included in the #PSM correlation plot.

### SasB activity assays

Michaelis-Menten characterization for SasB was carried out in a buffer containing 30 mM Tris pH 7.8, 150 mM KCl, 120 mM NaCl, 15 mM MgCl_2_, 1 mM TCEP, and 1mM pppGpp as an activator. ATP was included at a constant concentration of 8 mM and GDP/dGDP was varied from 1, 2, 5 and 10 mM. Final SasB concentration (counted as monomer) was 0.5 μM for ppGpp synthesis and 50 μM for pp(dG)pp synthesis and reactions were stopped after 90 seconds and 3 hours, respectively, by diluting 5μL into 1 mL 10 mM acetic acid. 200 μL diluted sample (1μL reaction) was resolved on MonoQ 5/50 using a gradient formed by Buffer A (5 mM Tris-HCl 8.0) and buffer B: (5mM Tris-HCl 8.0 and 1mM NaCl) with percentage of buffer B increasing from 20% to 40% over 10 mL. The peak corresponding to the product in the 254-nm chromatogram was quantified using the Evaluation module in the UNICORN 7.8 software. The extinction coefficient input was 0.013 and the “Amount” output was in units of mM from the original reaction.

### ppGpp, pp(dG)pp, ppApp and pp(dA)pp hydrolysis assays

All hydrolysis assays were performed in 96-well plates at 30 °C in a BioTek Synergy H1 microplate reader (Agilent). Production of GDP, dGDP, or dADP was coupled to ADP production using the activity of *B. subtilis* nucleoside diphosphate kinase (Ndk), and ADP production was quantitatively coupled to the stoichiometric consumption of NADH via the activity of excess pyruvate kinase (PK, type-III from rabbit muscle, Sigma) and lactate dehydrogenase (LDH, from rabbit muscle, Roche). Total reactions are as follows:

ppGpp + PEP + NADH + H^+^→ PPi + GTP + lactate + NAD^+^

pp(dG)pp + PEP + NADH + H^+^→ PPi + dGTP + lactate + NAD^+^

ppApp + PEP + NADH + H^+^→ PPi + ATP + lactate + NAD^+^

pp(dA)pp + PEP + NADH + H^+^→ PPi + dATP + lactate + NAD^+^

The volume of each reaction was 100 μL unless noted otherwise. Absorbance of each reaction was monitored for the absorbance at 340 nm (A340) every 1 minute. Concentrated stocks were prepared for reaction buffer, substrates, enzymes and additives and all reactions were assembled by first mixing all components except the substrate in a separate 96-well plate and pre-warmed to 30°C. Finally, the substrate was simultaneously transferred to multiple wells at t = 0 to initiate the reaction. Each reaction contained 50 mM HEPES-Na pH 7.5, 70 mM NaCl, 50 mM KCl, 10 mM NH_4_Cl, 0.1 mM TCEP, 50 μM ATP, 500 μM NADH, 7.5 mM PEP, 10 U/mL PK, 10 U/mL LDH, 1 μM B. *subtilis* Ndk, and appropriate concentrations of enzyme, substrate, divalent metal salt and/or EDTA as listed below.

- Michaelis-Menten kinetics (Figures 3C and D): reactions contained 5 mM MgSO_4_. Substrates were in the form of 1:1 mixture with MgSO_4_ at indicated final concentration to keep the free Mg^2+^ concentration constant. The final concentration of SpoT*^Ab^* was 100 nM (WT or Y51F mutant) for ppGpp hydrolysis and 1 μM for pp(dG)pp hydrolysis.
- Hydrolysis of ppApp vs pp(dA)pp and ppGpp vs pp(dG)pp (Figures 3F and G): reactions contained 5 mM MgSO_4_ and 500 μM substrate. Final concentration of enzymes was 100 nM for Aph1*^Bc^*, 1 μM for SAH*^Mex^*, 200 nM for MESH1 and 100 nM SpoT*^Ab^* when hydrolyzing the ribonucleotide substrate and 10-fold higher when hydrolyzing the 2’-deoxyribonucleotide substrate.
- Single metal cation (Mg^2+^ or Mn^2+^) titration reactions (Figures 4D and 5B) contained 500 μM ppGpp or pp(dG)pp. SpoT*^Ab^* was used at 30 nM for ppGpp hydrolysis and 1 μM for pp(dG)pp hydrolysis. MgSO_4_ or MnCl_2_ was included at final, total concentrations of 0.5, 0.75, 1, 1.5, 2, 3.5, 5, and 7.5 mM. When plotting for free metal concentration, 0.5 mM (1 equivalence to ppGpp) was subtracted from the total concentration.
- EDTA-buffered Mn^2+^ titration (Figure 4G): reactions contained 100 nM SpoT*^Ab^*, 500 μM ppGpp, 5 mM (black trace) or 30 mM (red trace) MgSO_4_, and 1 mM EDTA. MnCl_2_ was included at 50, 200, 400, 600, 800 and 900 μM (black symbols) or 50, 200, 400, 600, 800 μM (red symbols). Simulation of free Mn^2+^ concentration at each titration point is described in the next section.
- Mg^2+^ titration of EDTA-stripped SpoT*^Ab^* (Figure 4H): reactions contained 100 nM SpoT*^Ab^*, 500 μM ppGpp and 1 mM EDTA. MgSO_4_ was included at 0, 5, 10, 20, 35 and 50 mM. When plotting for free metal concentration, 1.5 mM (1 eq to ppGpp and EDTA) was subtracted from the total concentration.
- Single-point additive effects (Figures 4E, 5C and 5D): reactions contained 10 mM MgCl_2_ and 500 μM ppGpp or pp(dG)pp. SpoT*^Ab^* was used at 100 nM for ppGpp hydrolysis and 1 μM for pp(dG)pp hydrolysis. As indicated, reactions also include 1 mM EDTA or 100 μM MnCl_2_, CoCl_2_, NiSO_4_ or ZnAc_2_.

For each reaction, a background reaction was assembled on the same plate in which the hydrolase stock is replaced by the hydrolase purification buffer without the enzyme. All reactions were assayed together with their corresponding background reaction. Linear regression of the A_340_ time course provided initial velocity for each reaction in the unit of min^-1^. Upon background subtraction, this initial velocity is converted to μM•min^-1^ by dividing the extinction coefficient of 1.94x10^-3^ μM^-1^ for 100 μL reactions and 3.88x10^-3^ μM^-1^ for 200μL reactions.

### Simulation of free Mn^2+^ in the presence of Mg^2+^, EDTA and ppGpp

We first consider the distribution of EDTA across the following forms: completely protonated EDTA (denoted as “4^-^”), monoprotonated EDTA (3^-^), diprotonated EDTA (2-), Mg-EDTA chelate (MgEDTA) and Mn-EDTA chelate (MnEDTA). More protonated forms are negligible at our reaction pH of 7.5 provided the strong acidity of triprotonated EDTA (pKa = 2.7). [Mg^2+^] and [Mn^2+^] are used as the free metal concentrations. The weight of each species can be expressed as:

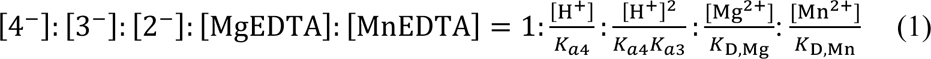

Constants *K*_a4_ = 10^-10.24^, *K*_a3_ = 10^-6.16^, *K*_D,Mg_ = 10^-8.79^ and *K*_D, Mn_ = 10^-13.89^ are available from literature (**ref**). Note that the weight of [4^-^], [3^-^] and [2^-^] is dependent only on pH, which is constant in our experiment. Because EDTA binds Mn^2+^ 5.1 orders of magnitude stronger than Mg^2+^, while ppGpp binds Mn^2+^ only 22-fold stronger than Mg^2+^ (5.5 μM vs 126 μM K_D_), any reactions containing **x** mM [Mg]_total_, 1 mM EDTA, 0.5 mM ppGpp and **y** (0.05 < y < 0.9) mM [Mn]_total_, the added Mn^2+^ is expected to displace Mg^2+^ from MgEDTA chelate rather than Mg-ppGpp chelate. Thus, [Mg^2+^] can be approximated to (**y**+**x**-1.5) mM, allowing us to calculate the weight for [MgEDTA] provided any [Mn]_total_ in this range. Additionally, [MnEDTA] proximates [Mn]_total_. At each [Mn]_total_, we solve [Mn^2+^] using equation (2).

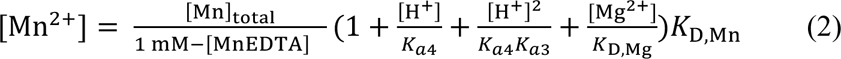

We then calculate the corresponding [4-] at each condition (see Table below). Note that [Mn] << [Mn]_T_ in agreement to our estimation that most Mn^2+^ forms chelate with EDTA by displacing Mg^2+^. To confirm that the quantity of Mn-ppGpp chelate is too little to affect our simulation, we used [Mg^2+^] at each condition and the *K*_D, Mg-ppGpp_ = 126 μM obtained in ITC experiment (SI Figure 3) to calculate free ppGpp concentration, [ppGpp]. We then used *K*_D, Mn-ppGpp_ = 5.5 μM to calculate the molar ratio between Mn-EDTA chelate and Mn-ppGpp chelate following equation (3). The results confirm that [MnppGpp] << [MnEDTA] in consistence with our basic assumption of the simulation process.

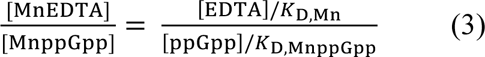

### Summary Table of simulation results. All units are M except the right most column

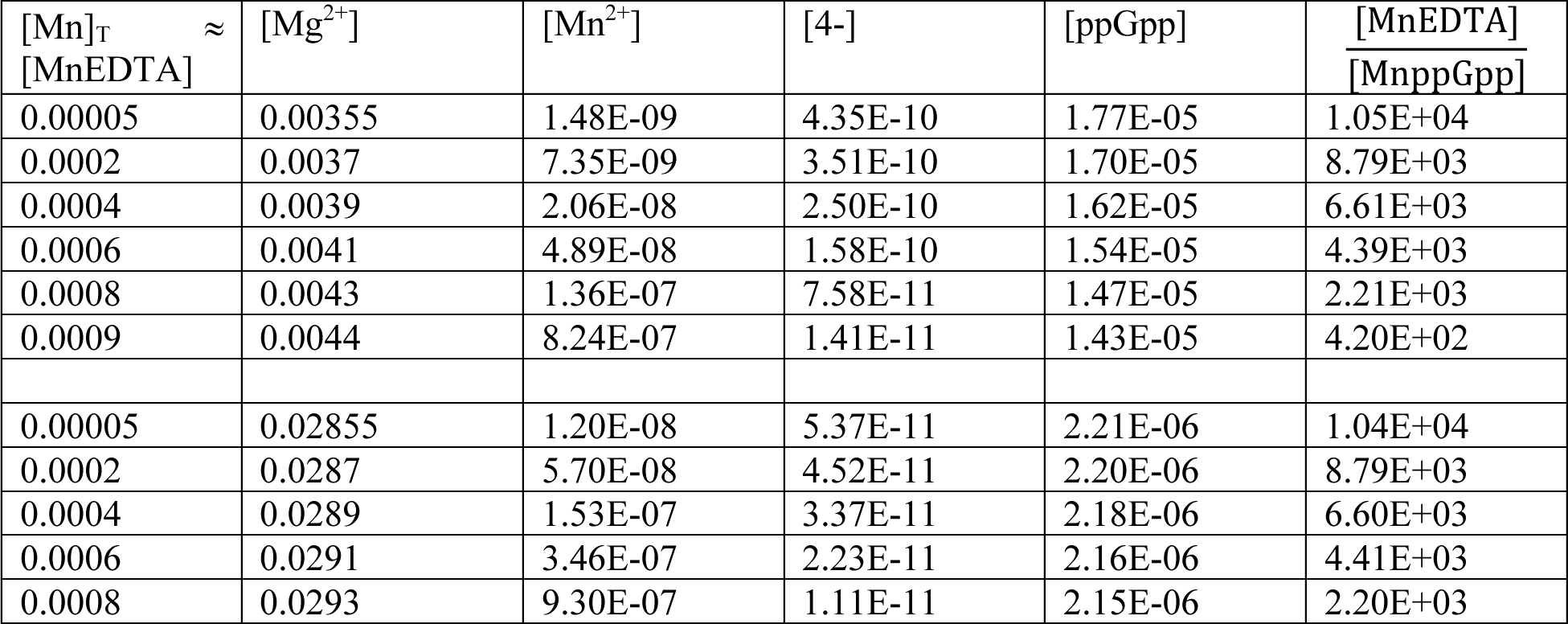

### Calculating *K_D_* for Mn^2+^ and Mg^2+^ from apparent *K_D_* values of Mn^2+^ at two different Mg^2+^ concentrations

We consider three states of M1 site: metal free, Mn^2+^-bound and Mg^2+^-bound and denote their concentrations as [M1], [M1Mn] and [M1Mg], respectively. The apparent *K_D_* of Mn^2+^ (*K*_D, App_) treats both metal-free and Mg^2+^-bound state “unbound” (because the latter is also almost inactive):

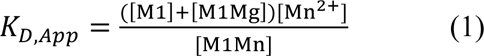

[M1] and [M1Mg] are related by *K*_D, Mg_ and free [Mg^2+^]:

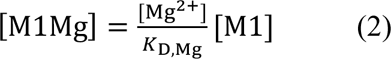

Thus,

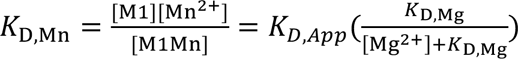

Using two sets of values:

*K*_D, App_ = 39 nM when [Mg^2+^] = 4 mM;

*K*_D, App_ = 116 nM when [Mg^2+^] = 29 mM;

We solved *K*_D, Mn_ = 27 nM and *K*_D, Mg_ = 8.6 mM.

### Multiple Sequence Alignment

All RSH sequences in Figure 3A were aligned using Clustal Omega ^5^ and YqeK*^Sp^* was aligned to SpoT*^Ab^* based on superimposition of their crystal structures (PDBID: 9WMY and 7QPR).

---

**Table S1.**
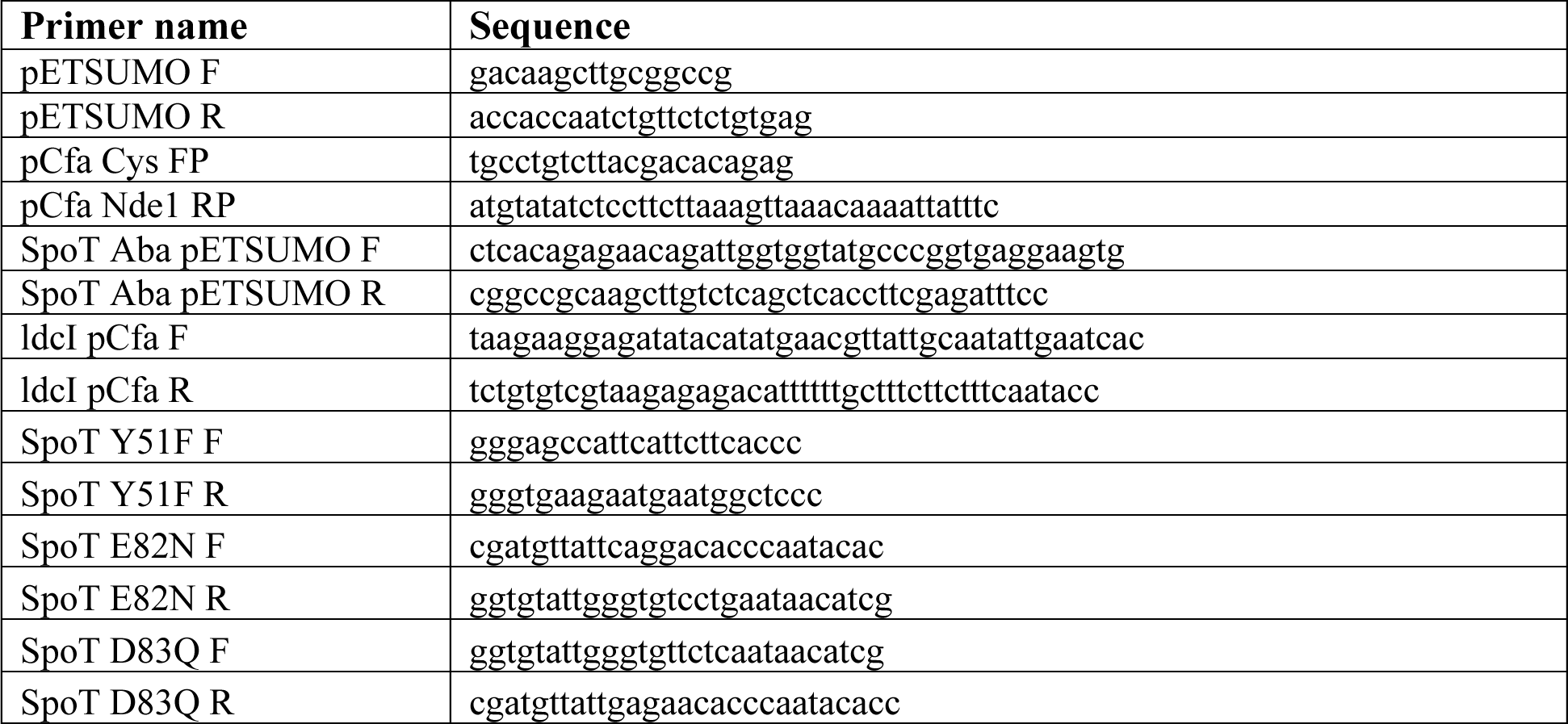
Primers.

**Table S2.** Plasmids.

| Plasmid Name | Relevant Genotype | Source |
| --- | --- | --- |
| P1-1 | P<sub>T7</sub> -His<sub>6</sub> -*yjbM*<sup>Bs</sup> (*sasB*) | Wang et al., 2019 |
| G74-6S | P<sub>T7</sub> -His<sub>6</sub> -*sumo-spoT*<sup>Aba</sup> | This study |
| G13-1 | P<sub>T7</sub> -His<sub>6</sub> -*speC*<sup>Eco</sup> | Wang et al., 2019 |
| G23-1Cfa | P<sub>T7</sub>-PurF<sup>Eco</sup>-Cfa-His<sub>6</sub> | Wang et al., 2019 |
| G24-1 | P<sub>T7</sub> -His<sub>6</sub> -GsK<sup>Eco</sup> | Wang et al., 2019 |
| G77-1Cfa | P<sub>T7</sub>-ldcI<sup>Eco</sup>-Cfa-His<sub>6</sub> | This study |
| G74-6S Y51F | P<sub>T7</sub> -His<sub>6</sub> -*sumo-spoT*<sup>Aba</sup> *Y51F* | This study |

**Table S6.** Strains.

| Strain # | Background | Plasmid | Source |
| --- | --- | --- | --- |
| 1 | MG1655 |  |  |
| 41 | BL21(DE3) | P1-1 | Wang et al. 2019 |
| 45 | BL21(DE3) | G74-6S | This study |
| 42 | BL21(DE3) | G13-1 | Wang et al. 2019 |
| 43 | BL21(DE3) | G23-1Cfa | Wang et al. 2019 |
| 44 | BL21(DE3) | G24-1S | Wang et al. 2020 |
| 45 | BL21(DE3) | G77-1Cfa | This study |
| 47 | BL21(DE3) | G74-6S Y51F | This study |

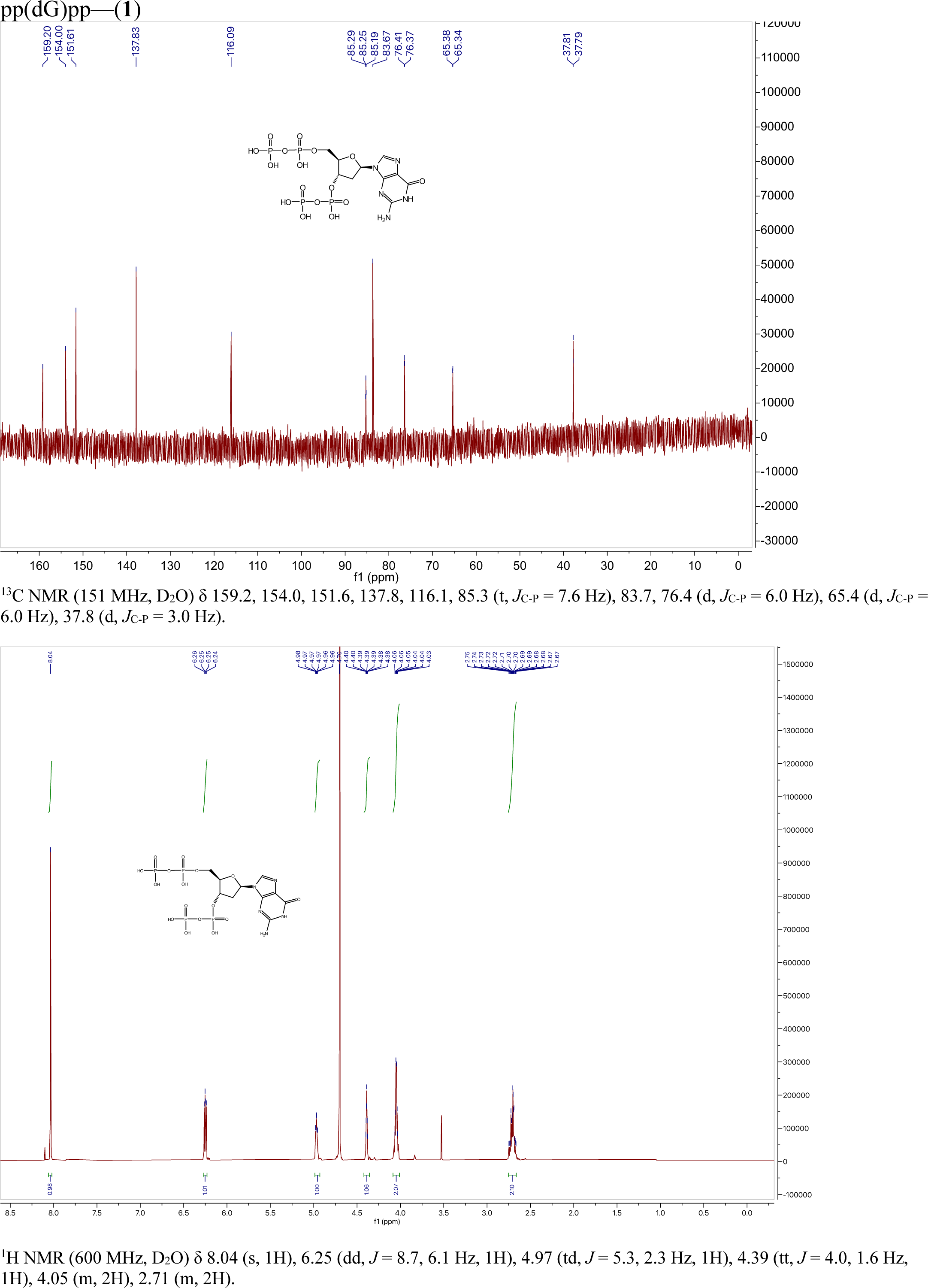

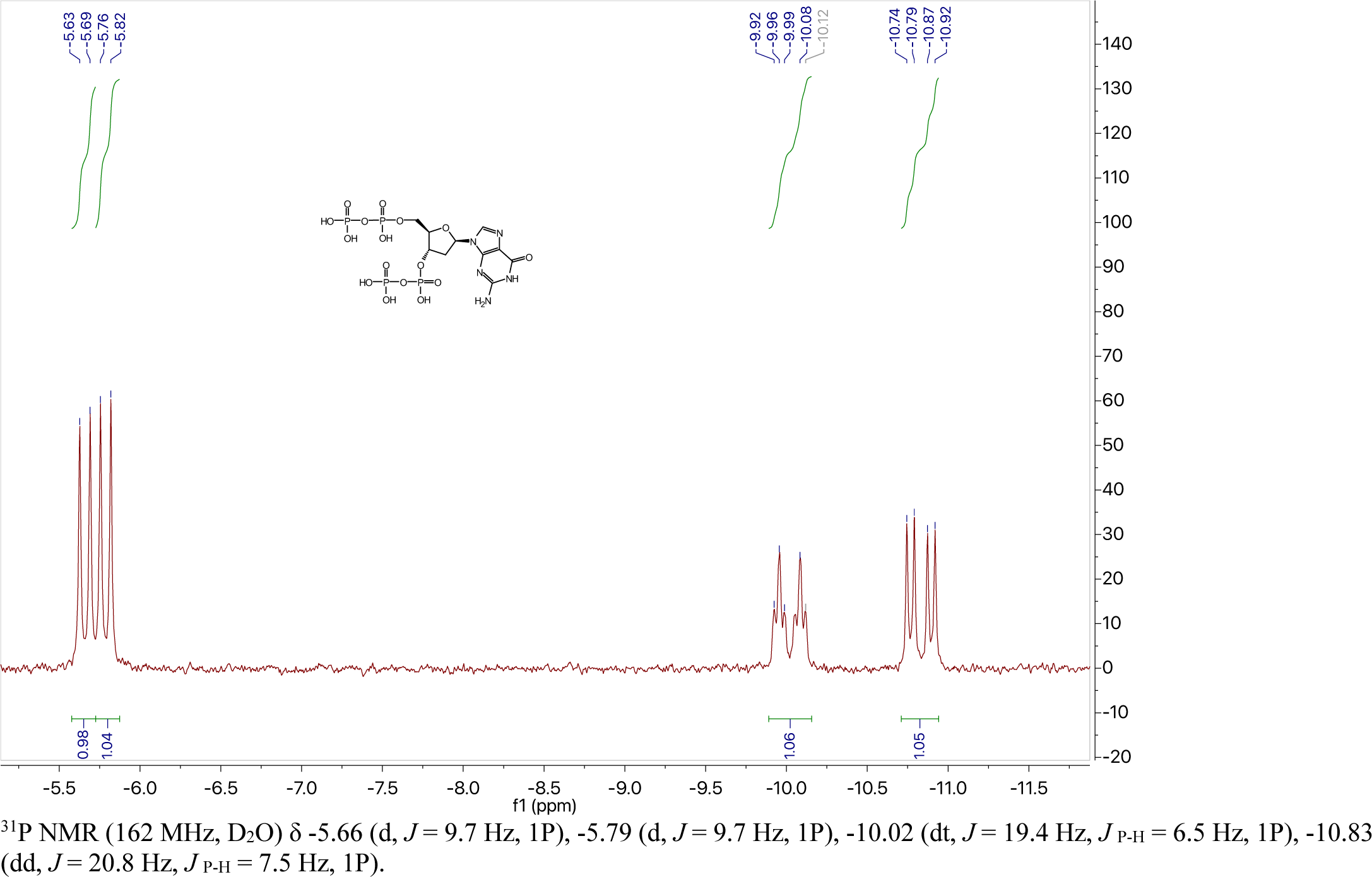

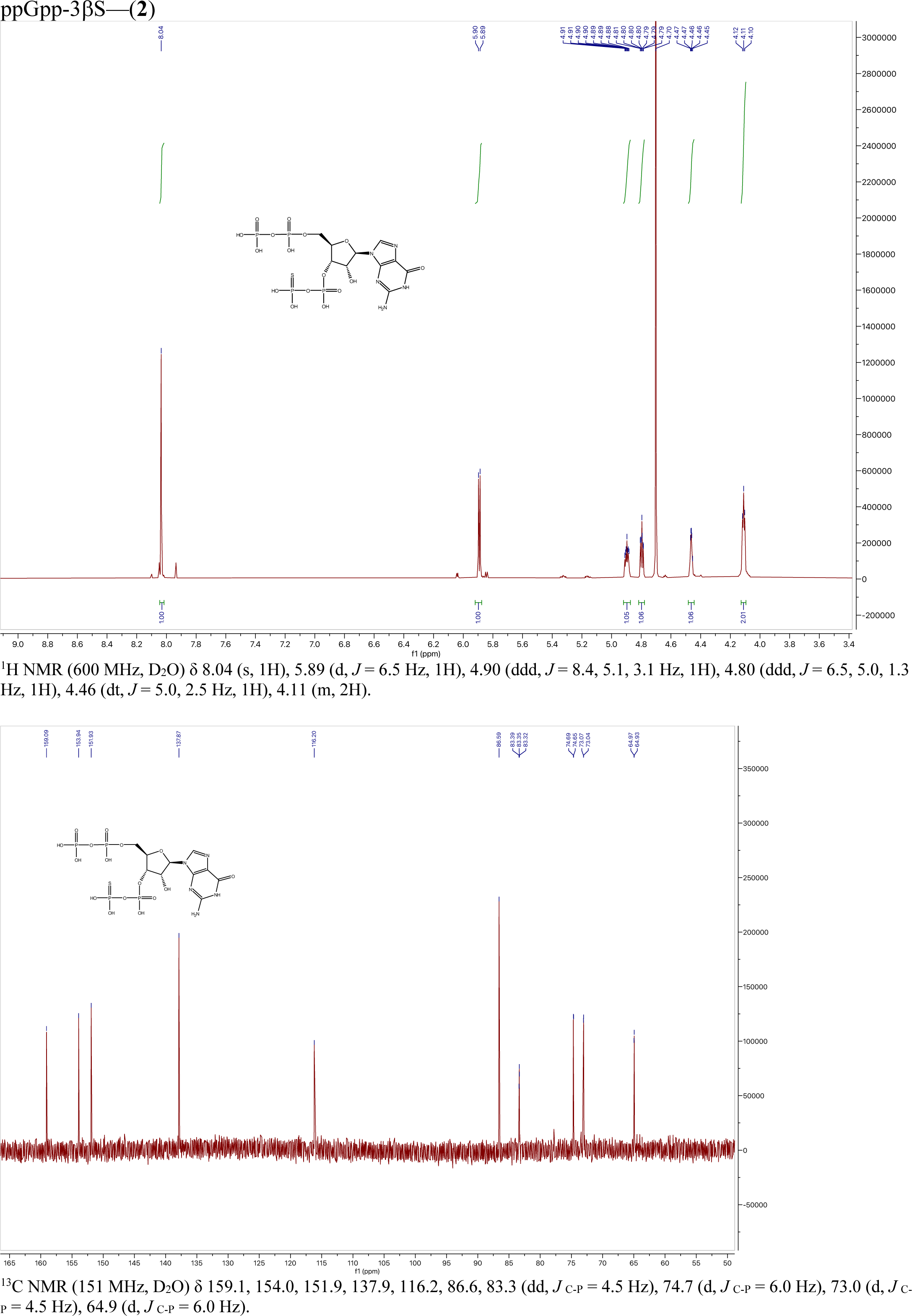

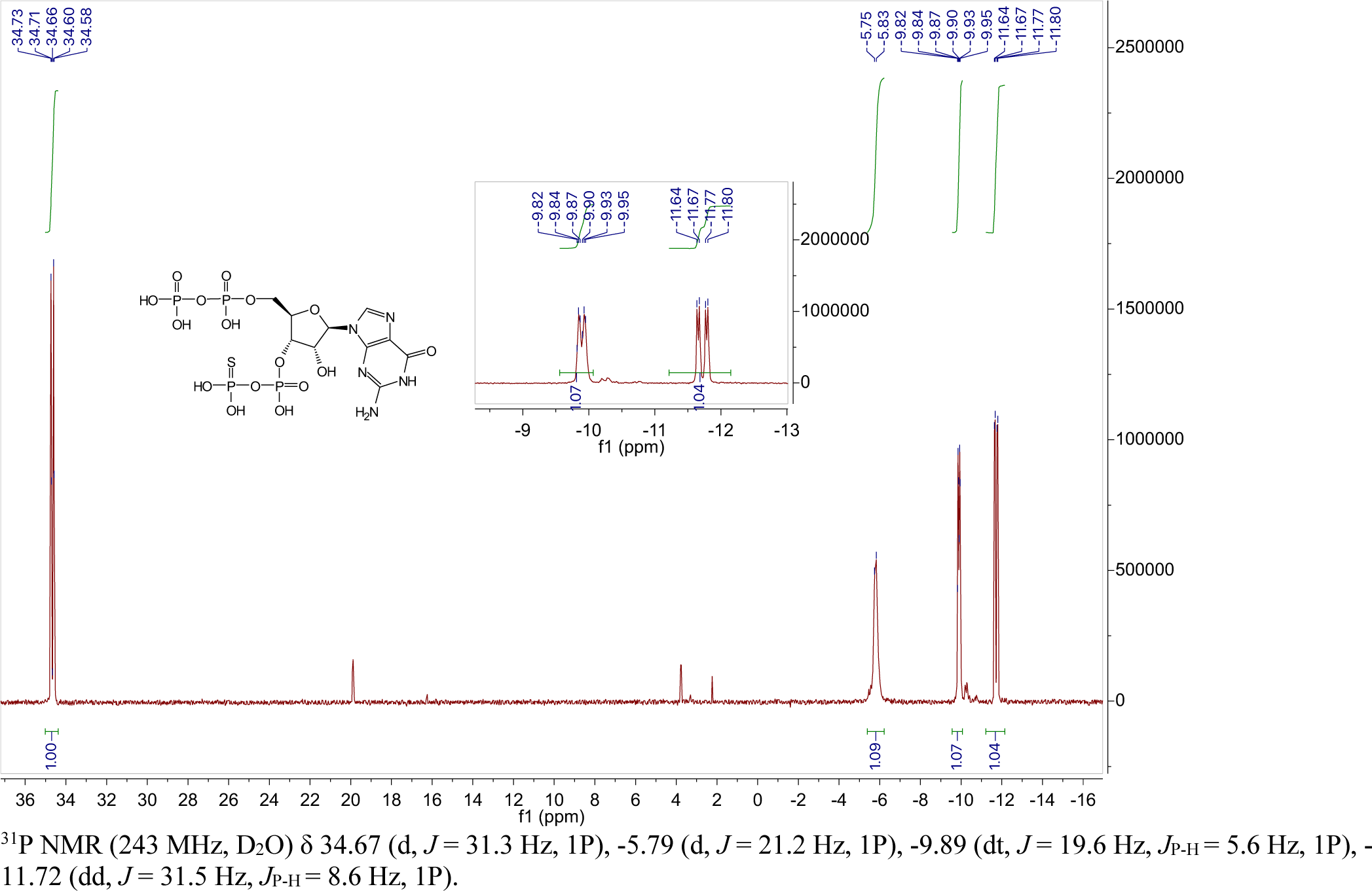

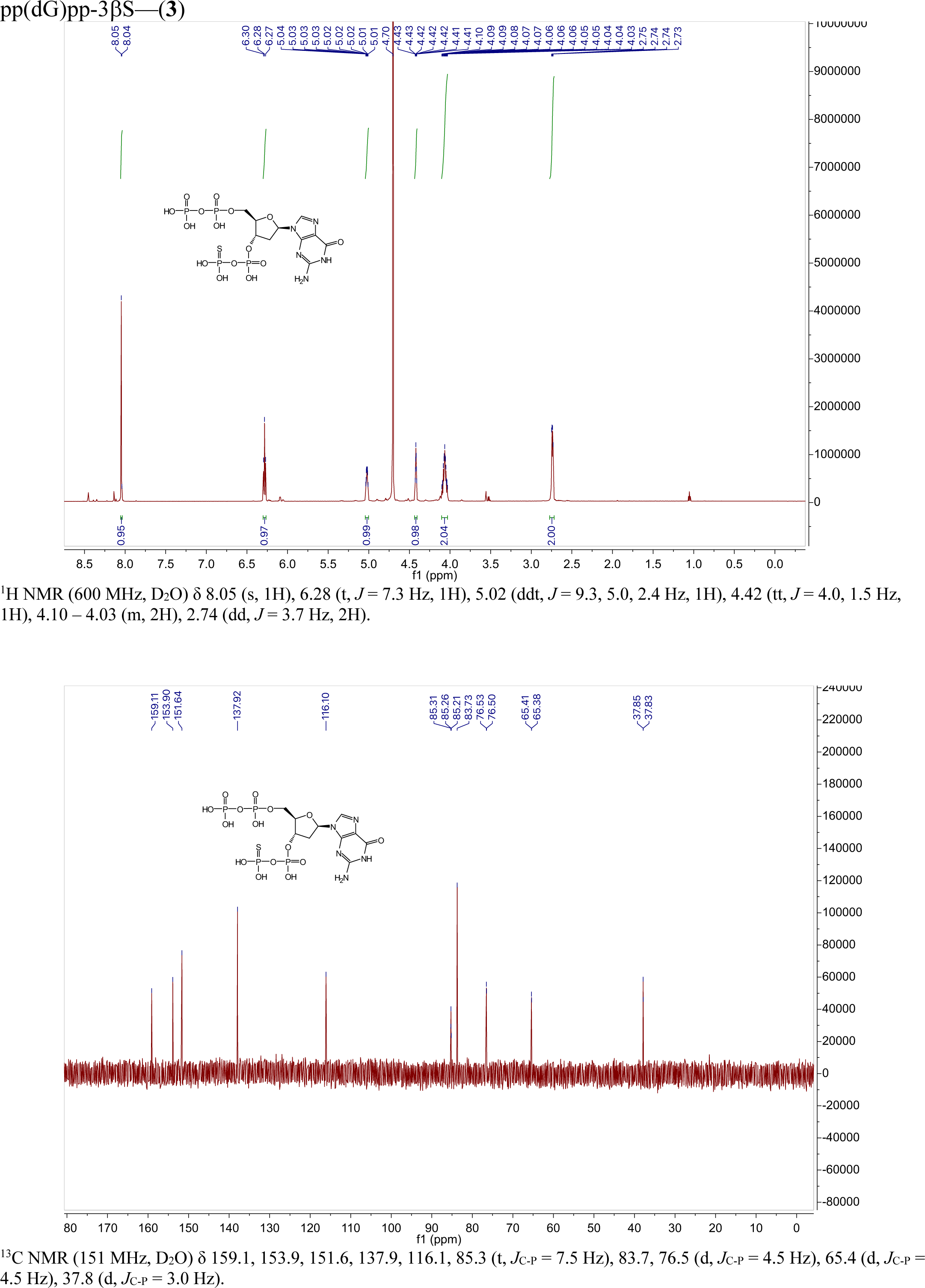

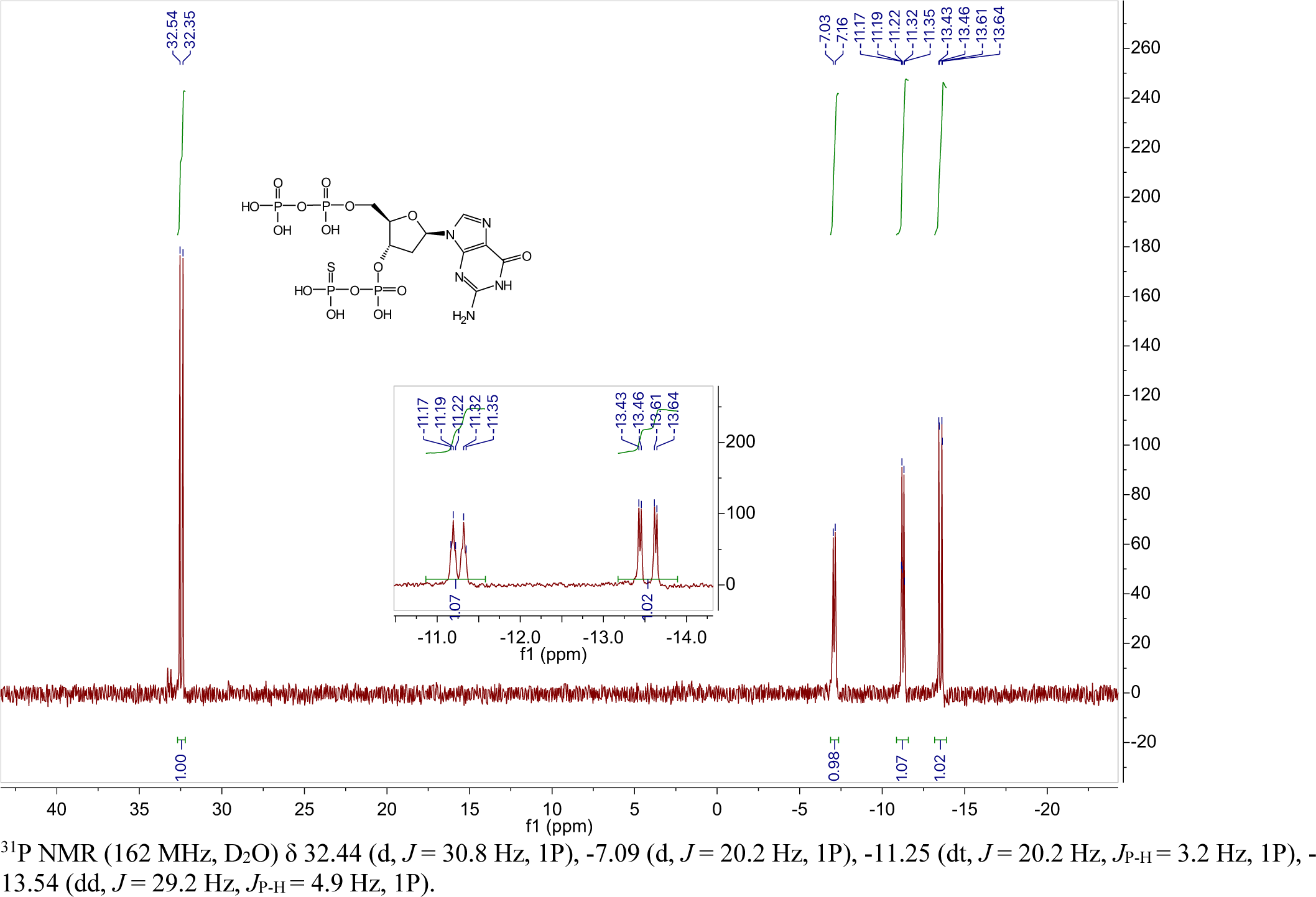

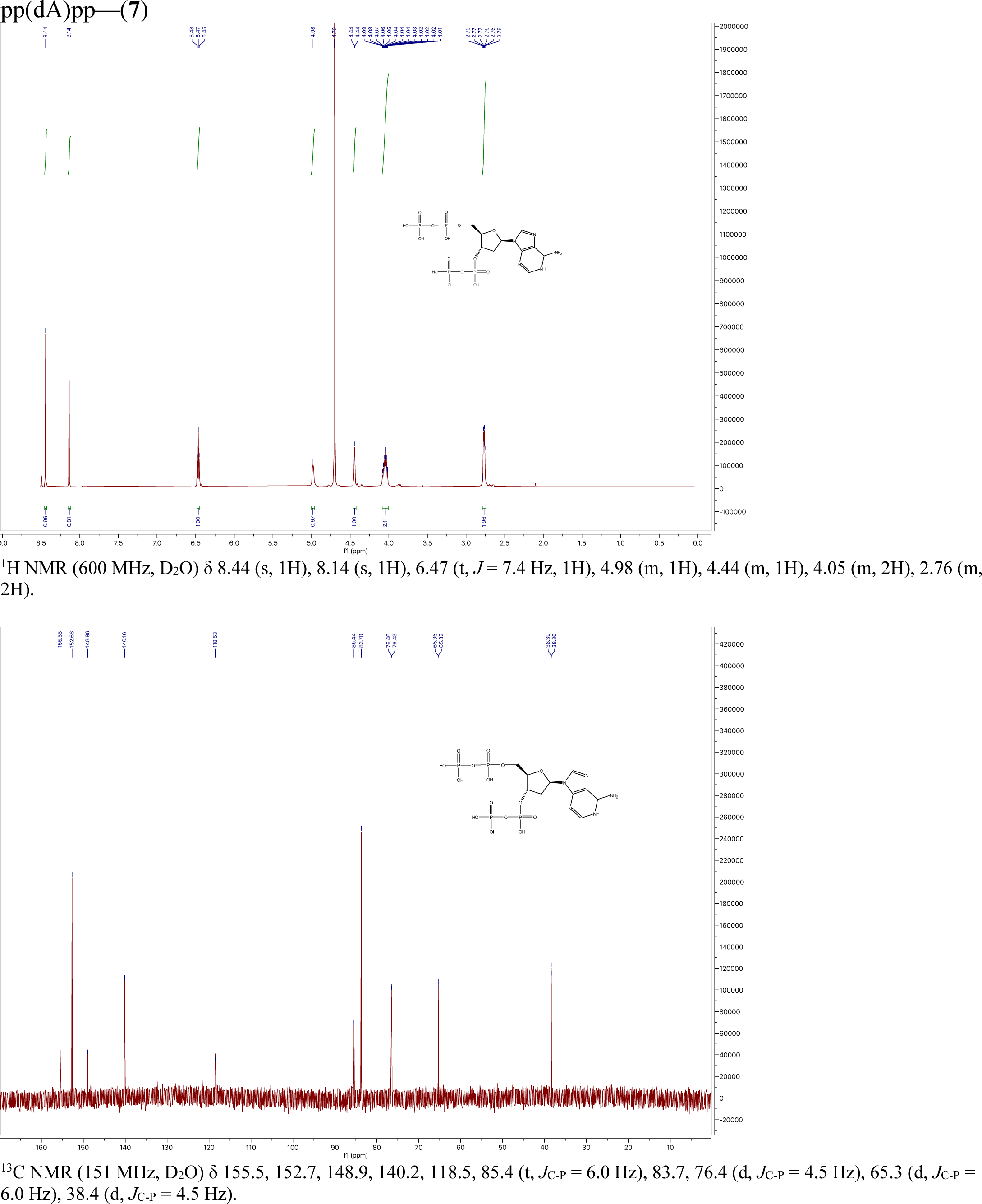

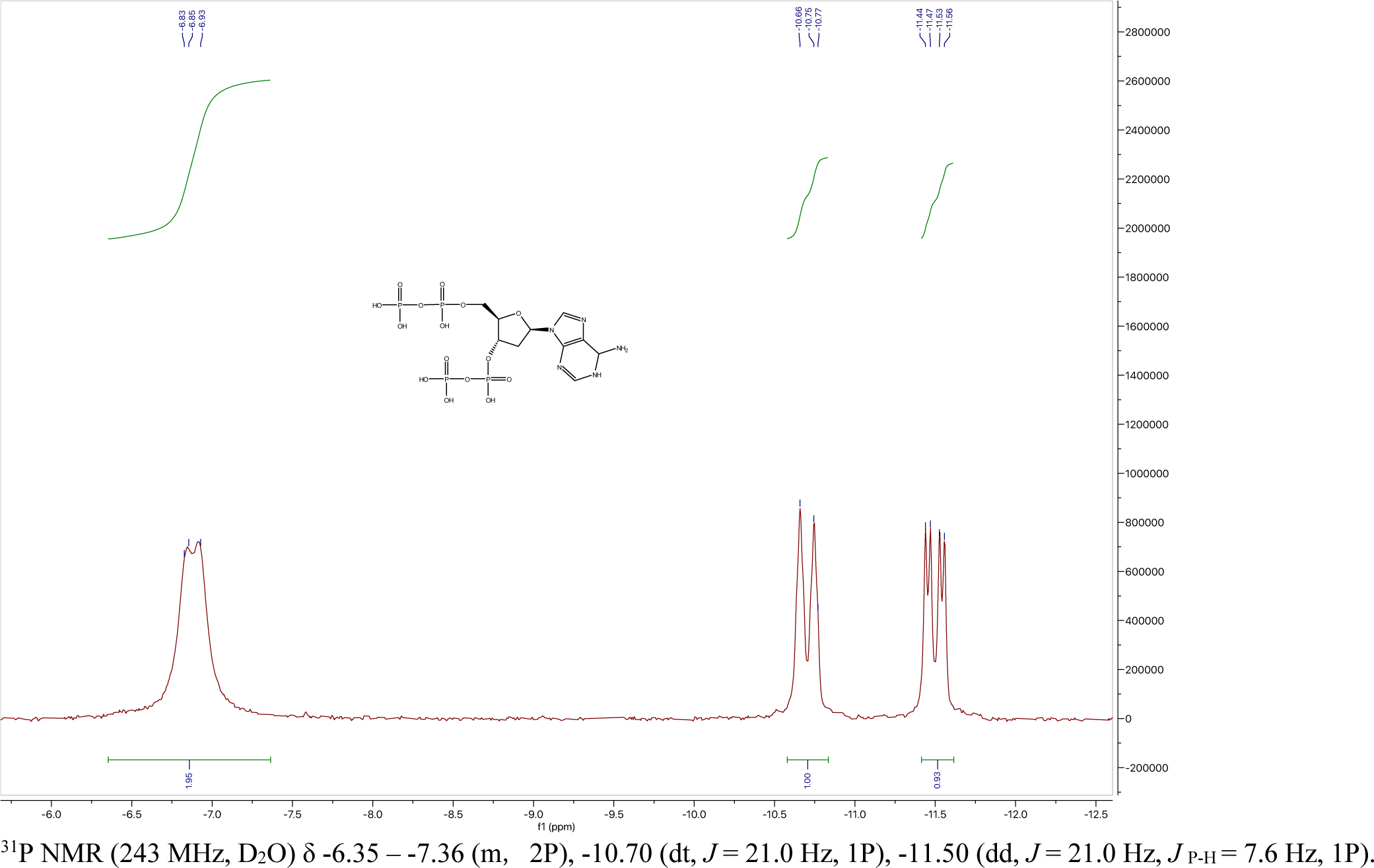

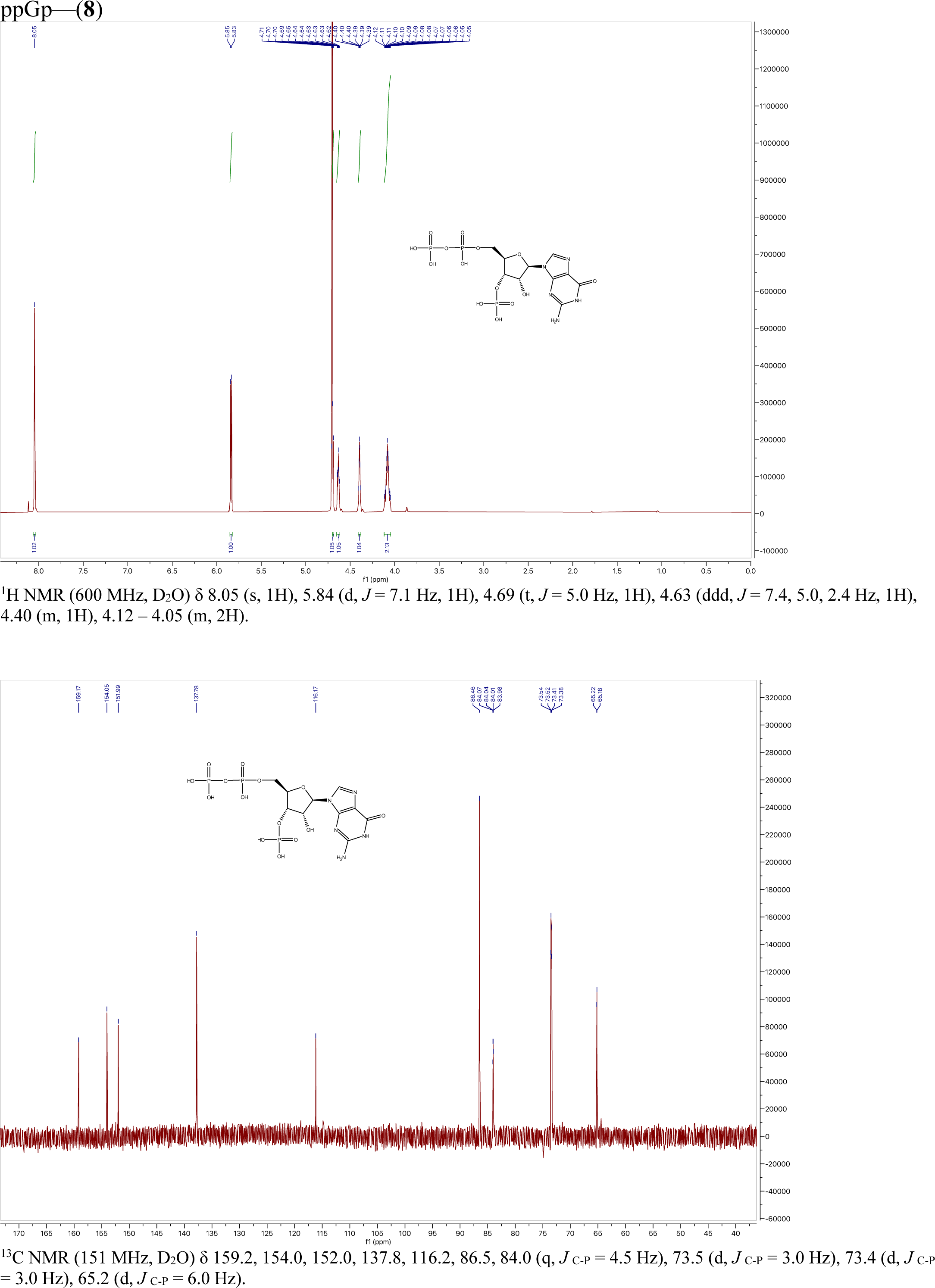

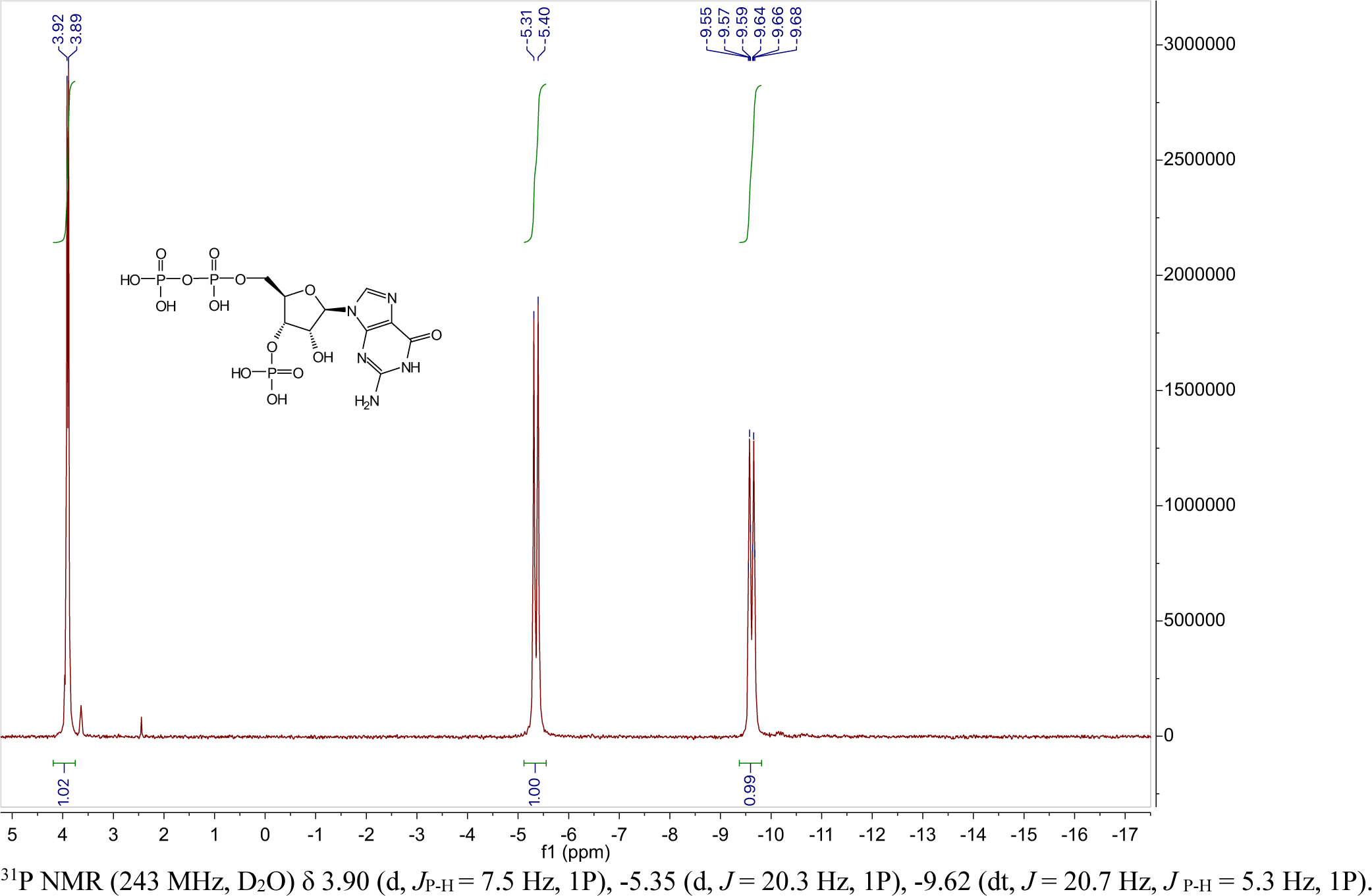

## Notes

### Competing Interest Statement

The authors have declared no competing interest.

